# Rendering Proteins Fluorescent Inconspicuously: Genetically Encoded 4-Cyanotryptophan Conserves Their Structure and Enables the Detection of Ligand Binding Sites

**DOI:** 10.1101/2024.09.18.613606

**Authors:** Haocheng Qianzhu, Elwy H. Abdelkader, Adarshi P. Welegedara, Edan Habel, Nathan Paul, Rebecca L. Frkic, Colin J. Jackson, Thomas Huber, Gottfried Otting

**Affiliations:** ARC Centre of Excellence for Innovations in Peptide & Protein Science, Research School of Chemistry Australian National University Canberra, ACT 2601, Australia; Research School of Chemistry, Australian National University, Canberra, ACT 2601, Australia

**Keywords:** cyanotryptophan, genetic encoding, fluorescence, ligand binding site, live cell imaging

## Abstract

Cyano-tryptophans (CN-Trp) are privileged multimodal reporters on protein structure. They are similar in size to the canonical amino acid tryptophan and some of them exhibit bright fluorescence which responds sensitively to changes in the environment. We selected aminoacyl-tRNA synthetases specific for 4-, 5-, 6-, and 7-CN-Trp for high-yield in vivo production of proteins with a single, site-specifically introduced nitrile label. The absorption maximum of 4-CN-Trp is distinct from Trp, allowing the selective excitation of its intense fluorescence. 4-CN-Trp features bright fluorescence in the visible range. Crystal structures of maltose binding protein demonstrate near-complete structural conservation when a native buried Trp residue is replaced by 4-CN-Trp. Besides presenting an inconspicuous tag for live cell microscopy, the high fluorescence of 4-CN-Trp enables measurements of subnanomolar ligand binding affinities in isotropic solution, as demonstrated by the complex between rapamycin and the peptidyl–prolyl isomerase FKBP12 furnished with a 4-CN-Trp residue in the substrate binding pocket. Furthermore, 4-CN-Trp residues positioned at different locations of a protein containing multiple tryptophan residues permits using fluorescence quenching experiments to detect the proximity of individual Trp residues to the binding site of aromatic ligands.

## Introduction

The amino acid tryptophan (Trp) endows proteins with a chromophore that has a high extinction coefficient and absorbs at relatively long wavelengths. These properties have made measurements of absorption at 280 nm the most commonly used method for determining protein concentrations.^[1]^ In addition, the Trp chromophore fluoresces with an emission maximum at about 350 nm, enabling the selective detection of Trp in a protein. As fluorescence presents the most sensitive detection technique in isotropic solution, the fluorescence of Trp has found countless applications, such as studies of protein folding and conformational transitions,^[2]^ and site-specific protein hydration.^[3]^ Of particular interest for drug discovery, Förster resonance energy transfer (FRET) and fluorescence quenching by aromatic ligands provide a straightforward way to identify binding partners and measure ligand binding affinities.^[4]^

For selective fluorescence labeling, either protein-selective or site-selective, the fluorescence marker must feature properties distinct from Trp. Several considerations apply. (i) For selective excitation and detection, the respective wavelengths of the fluorescence marker should be longer than those of Trp. Fluorescence in the visible range is highly desirable for protein tracking and microscopy. (ii) The fluorophore should be bright. (iii) To minimize structural perturbations in the protein, the fluorescence marker should be an a-amino acid with a small side chain. (iv) The marker should be easy to install. Genetic encoding of a non-canonical amino acid presents an elegant way for quantitative installation at a specific site.

Organic dye molecules typically achieve greater absorption wavelengths with an increased number of conjugated double bonds. Therefore, it is difficult to construct organic fluorophores smaller than the indole group of Trp. The non-canonical a-amino acids available for fluorescence emission in the visible range and shown to be suitable for installation in proteins in vivo by genetic encoding are based on coumarin,^[5]^ DansylA,^[6]^ 6-acetylnaphthalen-2-amine,^[7]^ and acridone.^[8]^ All these chemical groups are significantly larger than the indole group of Trp and prone to perturbing the structure of a target protein when installed in the hydrophobic core, while installation on the protein surface may significantly alter the interactions with other molecules.

To address these concerns, we developed a genetic encoding system for 4-cyanotryptophan (4-CN-Trp; Figure 1a). The fluorescence of 4-CN-Trp can be probed and detected independent of Trp, as the absorption maximum is at long wavelengths (about 320 nm), and the emission maximum is in the visible range (Figure 1b; about 425 nm in water, 395 nm in tetrahydrofurane).^[9]^ 4-CN-Trp differs from Trp by only two atoms, making it the smallest fluorescent amino acid known to emit in the visible range regardless of the polarity and H-bond donating capability of the environment.

**Figure 1.**
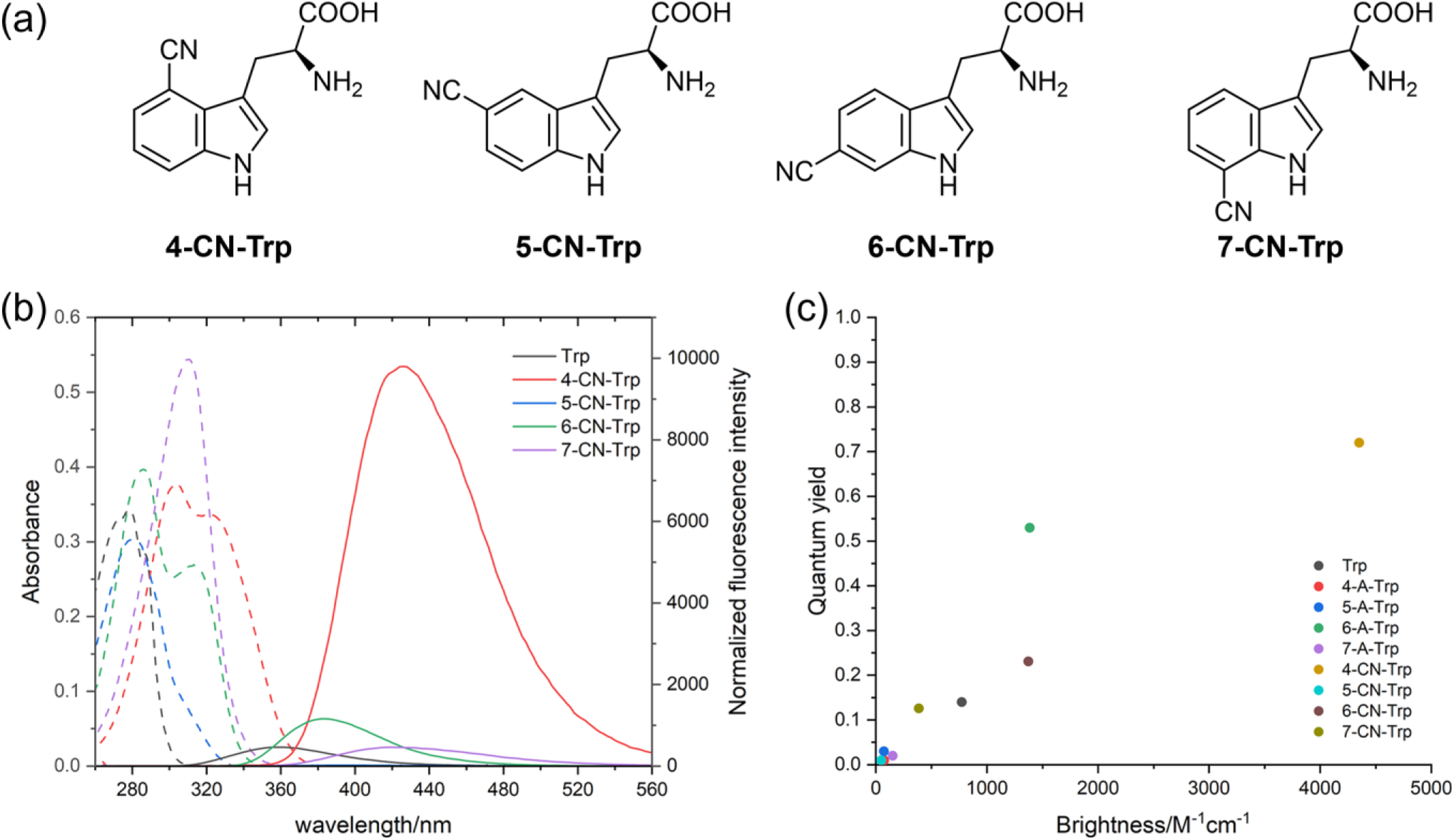
Cyanotryptophan isomers and their fluorescence spectra. (a) Chemical structures of the 4-, 5-, 6-, and 7-cyanotryptophan isomers. (b) Absorption (dashed lines) and emission (solid lines) spectra of 60 mM solutions of tryptophan and cyanotryptophan isomers in phosphate-buffered saline (PBS) buffer. The fluorescence emission spectra were acquired with irradiation at the wavelength of the absorption maximum of each amino acid (Table S1) and the fluorescence intensities were scaled by the absorbance at the respective excitation wavelength. (c) Quantum yield versus brightness for the different cyanotryptophans (CN-Trp) and comparison with tryptophan and azatryptophans (aza-Trp). Quantum yields were determined following the protocol of Würth et al.,^[22]^ utilizing tryptophan (λ_ex_ = 280 nm) or quinine sulfate (λ_ex_ = 305 nm) as reference standard. The brightness was calculated as the product of the extinction coefficient and quantum yield.

Besides allowing selective fluorescence excitation, 4-CN-Trp also offers much more attractive fluorescent properties than Trp. Trp fluorescence is complicated by featuring three major lifetimes (ranging from 0.5 ns to 5 ns),^[2]^ and the quantum yield is modest (about 0.15 in water).^[10]^ 4-CN-Trp fluorescence is much brighter by virtue of a higher absorption coefficient and a higher quantum yield (> 0.8 in water, > 0.4 in aprotic solvents). Finally, its fluorescence contains a long-lived component (> 12 ns) and it is more photostable than Trp, which is important for fluorescence microscopy.^[11]^

The fluorescence properties of 4-CN-Trp are also superior to the cyanotryptophan isomers with the cyano group in positions 5, 6, or 7 of the indole ring (5-CN-Trp, 6-CN-Trp, and 7-CN-Trp, respectively, Figure 1c). The quantum yield of 5-CN-Trp is very low in water (about 0.01). It is greater in 1,4-dioxane (0.11), but the emission maximum shifts to a much shorter wavelength in 1,4-dioxane (361 nm) compared with ca. 391 nm in water.^[12]^ 5-CN-Trp thus is a highly sensitive reporter of exposure to H_2_O, but its fluorescence can be difficult to separate from Trp because their absorption spectra overlap significantly and the brightness of 5-CN-Trp in water is low. 6-CN-Trp shows two absorption maxima, with the more intense absorption overlapping with that of Trp. The emission maximum of 6-CN-Trp is at about 370 nm. The fluorescence emission of 7-CN-Trp is relatively weak but extends beyond 420 nm in water (Figure 1c).

Excitation at 310 nm allows discrimination against Trp, but the fluorescence emission peak is highly dependent on solvent exposure and below 380 nm in methanol.^[13]^

Recently, genetic encoding of 5-, 6-, and 7-CN-Trp has been reported. Genetic encoding of non-canonical amino acids (ncAAs) ensures facile site-specific installation during in vivo protein synthesis in response to an amber stop codon. In the case of 5-CN-Trp, a mutant aminoacyl-tRNA synthetase (RS) based on the tyrosyl-RS system from *Methanocaldococcus jannaschii* has been shown to deliver high yields of two different proteins.^[14]^ 6-CN-Trp and 7-CN-Trp have been shown to be recognized by a chimeric pyrrolysyl-RS, but their installation in a wild-type GFP reporter proceeded in modest yields only.^[15]^

The cyano group possesses a polarity between that of a methylene and an amide group, and therefore can be accommodated both at buried and solvent-exposed sites with little energetic penalty.^[16]^ Despite the small structural difference between CN-Trp and Trp, the different CN-Trp isomers are not readily recognized by the wild-type *E. coli* tryptophanyl-tRNA synthetase.^[17]^ This is advantageous, as cyanotryptophans cause no damaging inhibition of cell growth even at relatively high concentrations, in contrast to, e.g., fluorotryptophans, which become toxic at concentrations greater than 2 mM. In particular, it has been noted that 4-CN-Trp is tolerated by *E. coli* at 0.25 mM concentrations without causing any significant growth inhibition.^[18]^ However, genetic encoding of 4-CN-Trp has remained elusive despite its unique attraction as a fluorescent probe.

In previous work, we developed systems for genetic encoding of 4-, 5-, 6-, and 7-fluorotryptophan^[19]^ and 7-azatryptophan,^[20]^ which are structurally even more similar to Trp than the CN-Trp isomers. None of these Trp analogues, however, allows selective excitation of their fluorescence without also exciting the fluorescence of canonical Trp, and none of them features an emission that is sufficiently resolved from the emission spectrum of Trp. The only potential exception is 6-azatryptophan, for which absorption and emission maxima have been reported at about 325 nm and 410 nm, respectively, with much greater variability depending on solvent exposure and side chain protonation than for 4-CN-Trp.^[21]^ Its fluorescence is considerably less bright (Figure 1c). A genetic encoding system for 6-azatryptophan has not been reported.

The present work describes efficient genetic encoding systems for 4-, 5-, 6-, and 7-cyanotryptophans based on the pyrrolysyl-tRNA synthetase of the methanogenic archaeon ISO4-G1 (G1PylRS),^[23]^ which has also been the basis for previously published genetic encoding systems of fluorotryptophan isomers and 7-azatryptophan.^[24,25]^ We describe the selection of requisite G1PylRS mutants from a library of mutant RS enzymes. We demonstrate protein production in high yield and purity. For 4-CN-Trp, we illustrate how the burial of the CN group in the hydrophobic core of a protein causes minimal structural perturbation in the protein structure, leaving the protein surface unchanged. Furthermore, we demonstrate the use of 4-CN-Trp for monitoring ligand binding with sub-nanomolar affinity in isotropic solution and for verifying ligand binding sites by fluorescence quenching.

## Results and Discussion

### Enzymatic Synthesis of the 7-CN-Trp Isomer from 7-Cyanoindole

While 4-, 5-, and 6-cyanotryptophan are readily available commercially, 7-CN-Trp is not. However, Arnold and coworkers described mutants of tryptophan synthase (TrpB) from *P. furiosus* and *T. maritima* that produce CN-Trp isomers from their corresponding cyanoindoles in good yield.^[24]^ We therefore expressed the *Tm*9D8^*^ mutant of TrpB with an N-terminal His6 tag in *E. coli* BL21(DE3) transfected with the pET-21(+) vector carrying the requisite gene (Twist Bioscience, USA). Following purification, 2.5 μM *Tm*9D8^*^ TrpB incubated at 55 °C for 72 h with 33 mM serine and 30 mM 7-cyanoindole yielded 7-CN-Trp in high yield and purity as confirmed by HPLC-MS and ^1^H NMR analysis (Figure S1).

### Fluorescence of CN-Trp Isomers

The absorption and emission wavelengths of the free amino acids in aqueous solution were investigated by measuring the fluorescence emission spectrum as a function of excitation wavelength. The results showed maximum emission of 4-CN-Trp at 432 nm following excitation at 320 nm (Figure S2, Table S1). Both in water and acetonitrile (Figure S3), 4-CN-Trp stands out among the CN-Trp isomers for combining long wavelengths of excitation and emission in the visible range with high brightness independent of the polarity and H-bonding capability of the environment, an excitation maximum beyond the range of canonical tryptophan.^[11]^

### Library Screening for 4-CN-Trp, 5-CN-Trp, 6-CN-Trp, and 7-CN-Trp tRNA Synthetases

The new RS enzymes were selected from the library of G1PylRS tRNA synthetases used previously for the genetic encoding of 7-azatryptophan and different fluorotryptophan isomers.^[19,20]^ *E. coli* DH10B cells were co-transformed with pBK-G1RS, which contains a library of G1PylRS mutants, and the selection plasmid pBAD-H6RFP, which encodes mCherry red fluorescent protein with an amber stop codon following an N-terminal His_6_ tag (His_6_-TAG-RFP). Relying on fluorescence-activated cell sorting (FACS), successive selection rounds changing between positive (with 1 mM ncAA provided) and negative selection (omitting any ncAA)^[25]^ increased the abundance of cells exhibiting the intended RS functionality across all four CN-Trp variants (Figures 1B and S4). After three selection rounds for 4-CN-Trp or five rounds for 5-CN-Trp, 6-CN-Trp, and 7-CN-Trp, multiple colonies of cells producing high RFP fluorescence were isolated and sequenced to identify 12 new G1PylRS mutants (Figure S5, Table S2). We refer to the top enzymes as 4CNWRS, 5CNWRS, 6CNWRS, and 7CNWRS. Samples of the expressed His6-TAG-RFP were purified and the specific incorporation of the various CN-Trp isomers was confirmed by high-resolution intact protein mass spectrometry (Figure S6).

### Protein Overexpression with Site-Specifically Installed CN-Trp

For large-scale production of proteins with CN-Trp residues, the genes of the G1PylRS enzyme mutants 4/5/6/7CNWRS were subcloned into the high-copy number plasmid pRSF, which also harbors the requisite amber suppressor tRNA.^[26]^ High efficiency and fidelity of amber read-through was again verified with the His6-TAG-RFP reporter gene carried by a pCDF plasmid under growth conditions with or without provision of the target CN-Trp (Figure 2). All four mutants showed no evidence of significant misincorporation of any of the canonical amino acids.

**Figure 2.**
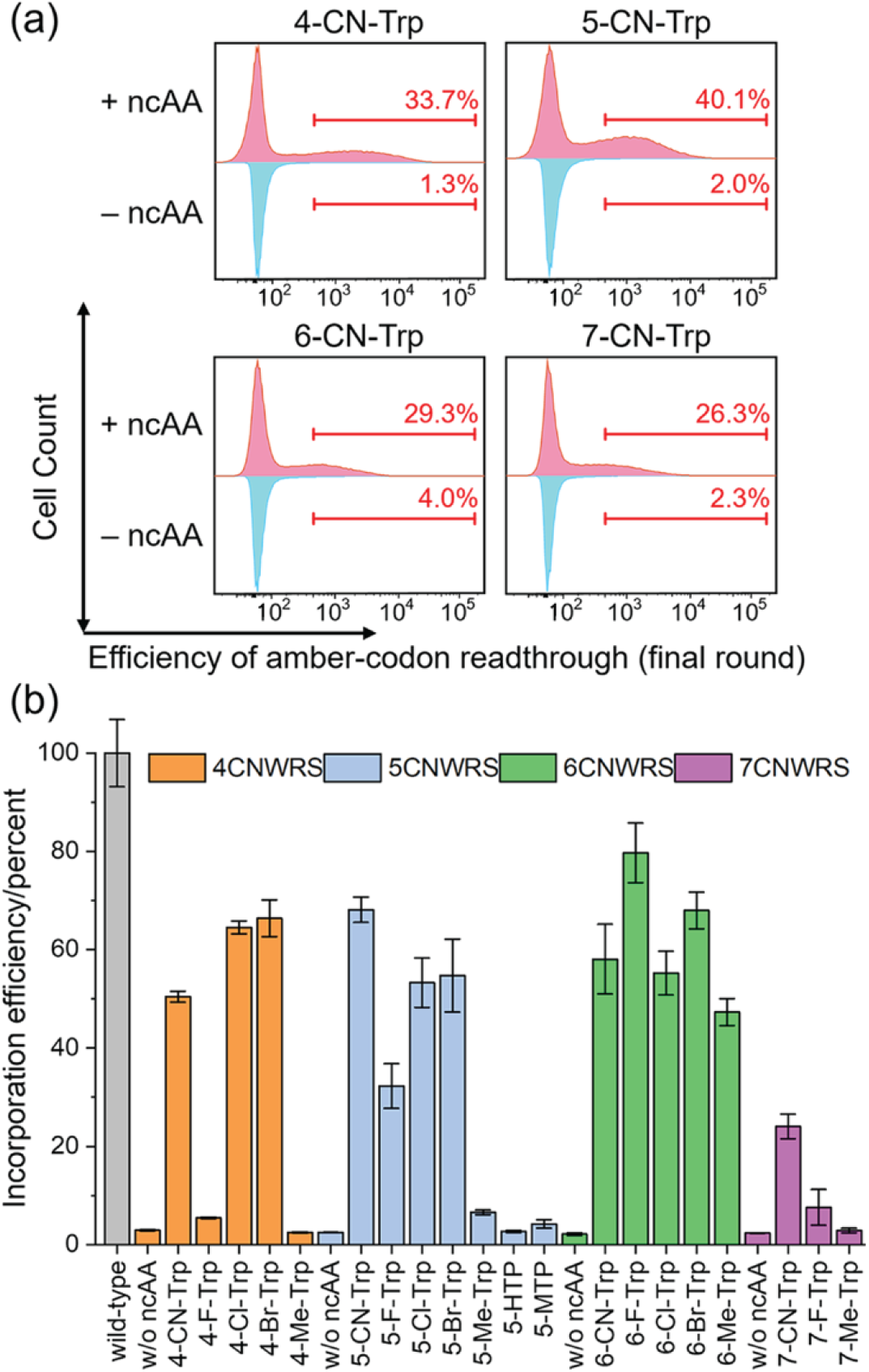
Identification of mutant G1PylRS enzymes recognizing different isomers of cyanotryptophan. (a) Chemical structures of the 4-, 5-, 6-, and 7-cyanotryptophan isomers and abbreviated names used in the present work. (b) Histograms of the final FACS selection rounds to identify G1PylRS enzymes active for 4-, 5-, 6-, and 7-cyanotryptophans, respectively. The horizontal axis represents the level of red fluorescence observed by expression of the mCherry red fluorescent protein (RFP) gene preceded by an amber stop codon. The vertical axis reports the cell count. The difference in RFP fluorescence intensity of cells grown with (red) and without (cyan) ncAA serves as an indicator of the presence of RS enzymes specific to the target cyanotryptophans in the gene pool. (c) Four selected tRNA synthetases for 4-, 5-, 6-, and 7-CN-tryptophan are polyspecific towards tryptophan isomers halogenated at the corresponding positions on the indole ring. Activity of these four enzymes to recognize ncAA variants as substrate was assessed by their efficiency to read through an amber-interrupted gene of RFP. Following cell growth in the presence of 1 mM ncAAs or without addition of any ncAA, the fluorescence intensity of expressed RFP was measured, normalized by OD, and compared to the cultures expressing wild-type RFP. Averages are shown and error bars indicate the standard deviation from three biological replicates.

As substrate polyspecificity has been reported for previously published aminoacyl-tRNA synthetases selected for ncAAs containing a cyano group,^[27]^ we also explored the capability of our newly identified enzymes to install other tryptophan-based ncAAs (Figure S7). Tryptophan analogues with halogen atoms in the same position as the cyano group in the CN-Trp isomers were incorporated into our test protein by the respective G1PylRS mutants (Figure 2b). In addition, 6-methyl-typtophan (6-Me-Trp) was recognized as the substrate of 6CNWRS with high efficiency.

Proteins of interest with 4-CN-Trp were overexpressed in *E. coli* B95.ΔAΔ*fabR* cells,^[28]^ which are devoid of the release factor RF1, transfected with the pRSF plasmid carrying the 4CNWRS gene. As in the experiments conducted to express RFP variants with different tryptophan analogues, the gene of the target protein with the amber stop codon was provided in the low-copy number plasmid pCDF and co-transfected together with the pRSF plasmid (Figures S8 and S9). Following expression with 4-CN-Trp supplied in 1 mM concentration, intact protein mass spectrometry of the purified proteins demonstrated successful incorporation of the ncAA without misincorporation (Figures 3, S10, and S11). Protein yields ranged between 7 mg and 20 mg for the maltose binding protein (MBP) and the ligand binding domain of the human pregnane X receptor (hPXR-LBD) and 63 mg for the peptidyl–prolyl *cis*–*trans* isomerase FKBP1A (FKBP12) per liter cell culture. The blue fluorescence is easily observed on SDS-PAGE without staining (Figure 3b) and readily detected even at low concentrations (Figure 3c). Good expression yields of full-length protein were obtained also with *E. coli* protein expression strains that contain the RF1 release factor (Figure S12).

**Figure 3.**
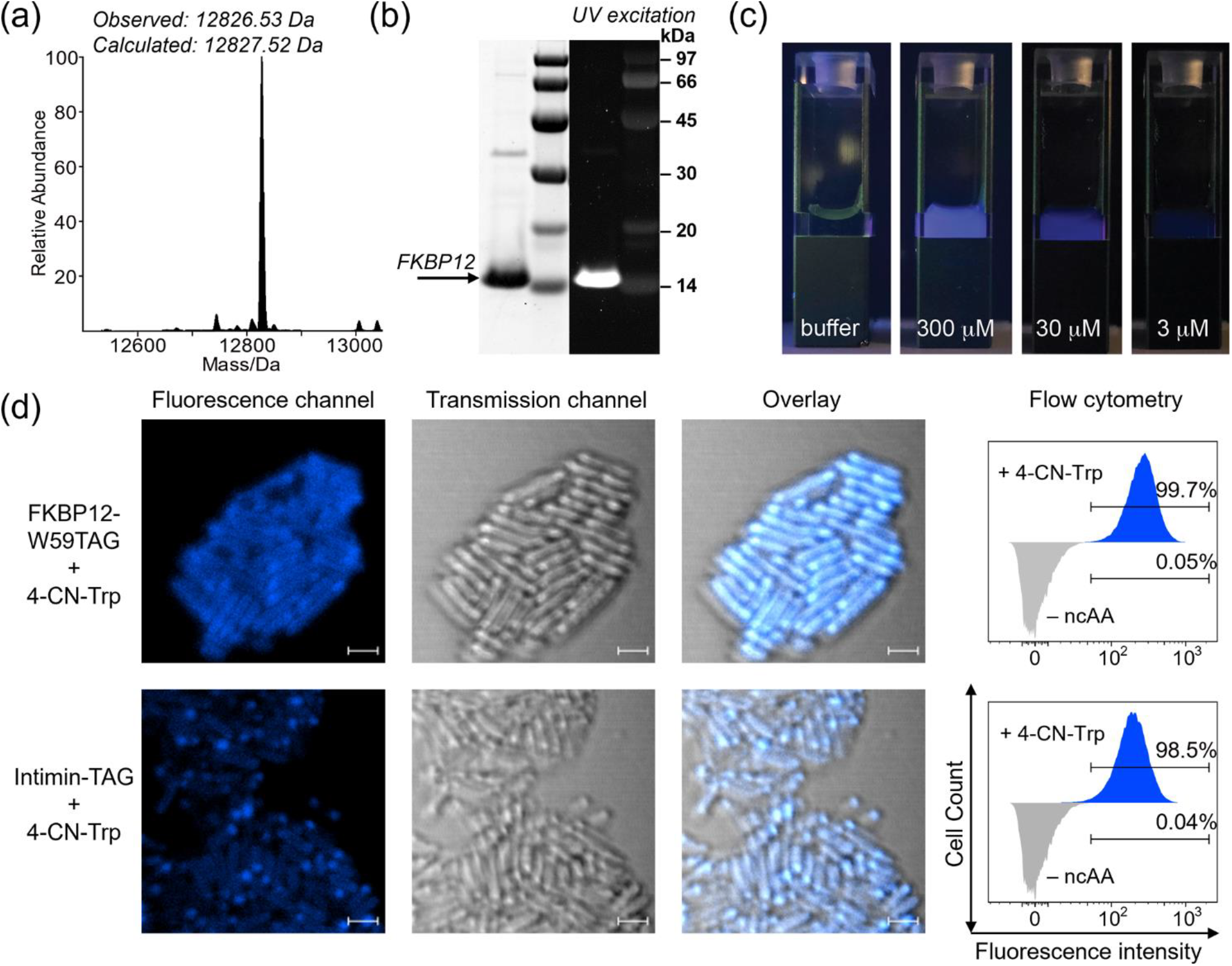
Blue fluorescence conferred to a protein by a single 4-CN-Trp residue. Trp59 in the protein FKBP12 was replaced by 4-CN-Trp (FKBP12-4CNW). (a) ESI intact protein mass spectrometry confirming the incorporation specificity. (b) The SDS-PAGE band of FKBP12 with 4-CN-Trp is readily detected by fluorescence without staining. The white-background image on the left shows the gel after staining with Coomassie Blue. The dark-background image on the right was taken before staining the gel, using a 302 nm UV transilluminator for excitation and a 590/110 nm emission filter for detection. (c) Photo showing the fluorescence of the protein in a 1 cm quartz cuvette at different concentrations in PBS buffer. The fluorescence was excited by irradiation at 320 nm and photographed with an ordinary mobile phone camera. (d) Confocal fluorescence microscopy images and flow cytometry analysis of live *E. coli* cells expressing either FKBP12-4CNW or displaying the intimin autotransporter with 4-CN-Trp incorporated at its C-terminus (scale bar = 2 μm). Fluorescence was excited using 355 nm lasers. The microscopy images were processed combining the emission signals below 503 nm. Flow cytometry analysis of the populations of fluorescent cells was performed with a 379/28 bandpass filter. In the positive sample (blue), almost all cells expressed FKBP12-4CNW. In the negative control (gray), where cells were cultured without supplying 4-CN-Trp, cells with similar fluorescence intensity are rare.

### Fluorescence Imaging of Live Cells with 4-CN-Trp

To explore the utility of 4-CN-Trp for live fluorescent imaging, we prepared *E. coli* cells expressing various amber-interrupted reporters: (1) FKBP12-4CNW which is a soluble intracellular protein, (2) an intimin autotransporter construct^[29]^ with an amber stop codon near the C-terminal end of the intimin passenger domain, Intimin-TAG, to display 4-CN-Trp on the cell surface, (3) a construct of the *E. coli* outer membrane protein X,^[30]^ OmpX-TAG, where 4-CN-Trp was inserted into an extracellular loop, and (4) an amber-interrupted nanobody gene fused with the pelB signal peptide, PelB-NB-F67TAG,^[31]^ for localization of a 4-CN-Trp-containing protein in the periplasmic space. Baseline controls were obtained by *E. coli* cells lacking the incorporation system but incubated with the same concentration of 4-CN-Trp. Negative control samples were cultured without 4-CN-Trp supplementation. After overnight expression and thoroughly washing the cells with PBS buffer, the bacteria were immobilized on agarose pads and imaged by confocal microscopy (Figures 3d, S13, and S14). In addition, washed cells were analyzed by flow cytometry to measure the fraction of cells containing fluorescently-tagged proteins. This ratio was nearly 100% (Figure 3d). Non-specific binding of free 4-CN-Trp was minimal (Figure S13). The fluorescence images revealed a uniform distribution of FKBP12-4CNW in the cytosol, whereas the membrane-bound and periplasmic proteins showed uneven spatial distributions.

### Site Dependence of 4-CN-Trp Fluorescence

The wavelengths of the fluorescence emission maxima of cyano-indoles and cyano-tryptophans are known to depend on the polarity and H-bonding capacity of the solvent (Figures S2 and S3),^[9]^ and this effect is also observed in the proteins containing 4-CN-Trp in positions of different solvent exposure. For example, one-by-one substitution of the eight Trp residues in the MBP for 4-CN-Trp delivered emission maxima in the range between 380 and 410 nm (Figure S15a). In 8 M urea, the emission maxima were all at 410 nm independent of the location of the 4-CN-Trp residue in the amino acid sequence (Figure S15b), demonstrating high solvent exposure in the unfolded protein.

Protein-dependent differences in wavelengths of the fluorescence emission maxima can also be detected in live cells. spectra of bacteria carrying proteins with 4-CN-Trp—whether intracellular, surface-displayed, or located in the periplasmic space—were recorded and compared (Figure S16). For example, the cells displaying 4-CN-Trp in the intimin-TAG and OmpX-TAG constructs exhibited fluorescence maxima at greater wavelengths (ca. 413 nm) than the cells expressing FKBP12-4CNW and PelB-NB-F67TAG (Figure S16). Conceivably, differences observed between purified and in vivo samples could be used to detect interactions privileged in the in vivo environment.

### Structural Conservation of MBP with 4-CN-Trp

To investigate the structural impact arising from the switch of a native Trp residue to a 4-CN-Trp residue, we determined the 3D structure of MBP with 4-CN-Trp in position 10 by X-ray crystallography. The 1.5 Å resolution structure revealed a conserved orientation of the 4-CN-Trp side chain despite its partial solvent exposure and the position of the cyano group, which points into the hydrophobic core of the protein. This indicates that the cyano group is readily accommodated at this site. Compared to the crystal structure of the wild-type protein crystallized in the same conditions, the χ_1_ and χ_2_ angles of 4-CN-Trp differ from the wild-type by less than 5° and the χ_1_ and χ_2_ angles of the nearby fully buried residue Ile60 by less than 13° (Figure 4). These structural adaptations are even smaller than those we previously observed in the hydrophobic core of a protein produced with leucine and valine analogues containing single hydrogen-to-fluorine substitutions in the methyl groups.^[32]^ Importantly, the surface of MBP is practically unaffected by the substitution of a single Trp residue for a 4-CN-Trp residue (Figure S17).

**Figure 4.**
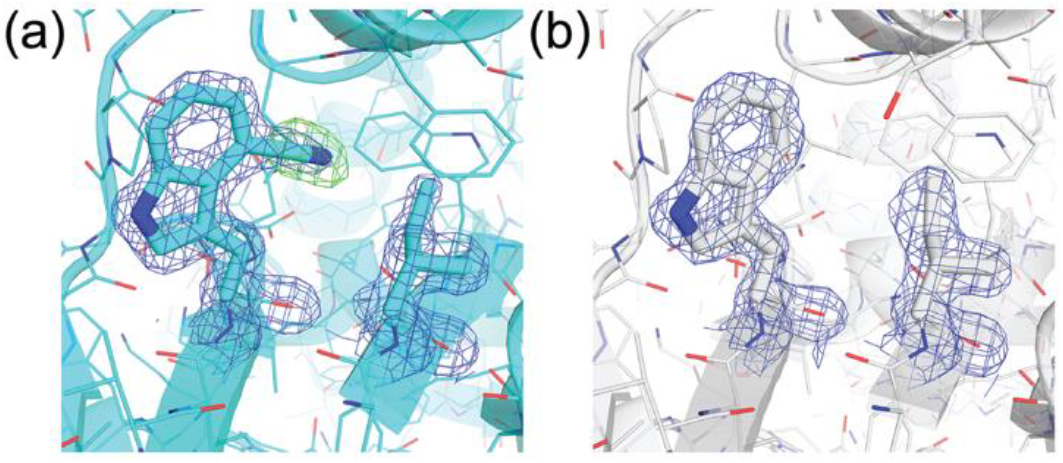
Comparison of the side chain conformations of Trp and 4-CN-Trp in position 10 of native and mutated MBP, illustrating the level of structural conservation. (a) MBP produced with a single 4-CN-Trp residue in position 10 (PDB ID 9CLC). 2mF_O_-DF_C_ densities are shown as a blue mesh contoured to 1.5 σ, highlighting the conformation of the 4-CN-Trp residue in position 10 and Ile60. Omit density (modeled without the atoms of the CN group) is shown in green mesh contoured to 3.0 σ. (b) Same as (a), but for wild-type MBP (PDB ID 9LCD).

### Ligand Binding Affinities in the Subnanomolar Range

Besides endowing proteins with bright intrinsic fluorescence, the installation of 4-CN-Trp in proteins delivers a sensitive site-specific probe of changes in the local chemical environment, which can be exploited for monitoring ligand binding events by direct fluorescence measurements in isotropic solution. For example, FKBP12 is known to bind rapamycin with a very low dissociation constant (*K*_d_ = 0.2 nM).^[33]^ The fluorescence emission spectrum (excitation wavelength 310 nm) of FKBP12-4CNW recorded at 1 μM concentration in PBS buffer responds sensitively to the presence of 20 μM rapamycin, with the maximum of the emission peak shifted from 379 nm to 371 nm after ligand binding, accompanied by about 14% decrease in fluorescence intensity (Figure 5a). Rapamycin does not contain aromatic moieties that could act as fluorescence quenchers, suggesting that the changes observed in the emission peak arise from rapamycin changing the solvent accessibility of the cyano group when the ligand binds. A control experiment monitoring the fluorescence of 4-CN-Trp instead of FKBP12-4CNW confirmed that rapamycin exerts a much lesser fluorescence quenching effect on the free amino acid (Figure S18). The decrease in fluorescence intensity upon ligand binding is still measurable at 4 nM protein concentration in a 1 cm cuvette (Figure S19, a and b). The reduction in fluorescence intensity observed in a titration experiment with rapamycin yielded a classical binding curve, from which a *K*_d_ value of 0.4 nM was derived by fitting the equation for two-state binding (Figures 4b and S19c).

**Figure 5.**
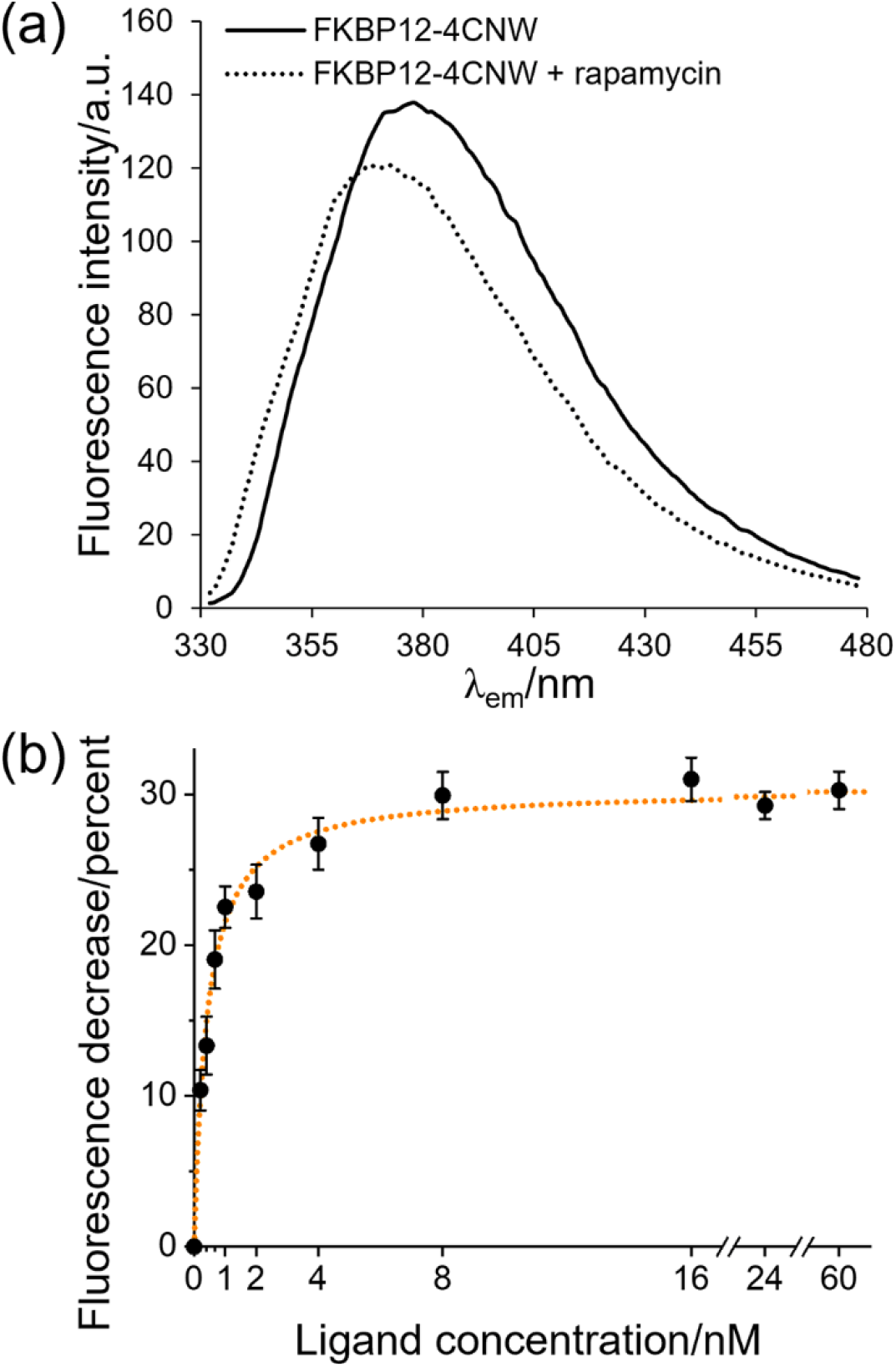
Detecting the binding of rapamycin to FKBP12-4CNW by fluorescence. (a) Fluorescence emission spectra of 1 μM FKBP12-4CNW. The spectra were recorded at room temperature in PBS buffer using an excitation wavelength of 310 nm. Addition of a 20-fold excess of rapamycin yields a blue shift of the emission maximum from 378 nm to 371 nm and a decrease in emission intensity. (b) Decrease in fluorescence upon ligand binding versus rapamycin concentration in a 4 nM solution of FKBP12-4CNW. The background fluorescence with rapamycin only in PBS buffer was subtracted from each spectrum and the decrease in fluorescence intensity plotted. The error bars indicate the standard deviation from triplicate measurements.

When the ligand dissociation constant, *K*_d_, is determined by monitoring changes in the protein upon titration with ligand, it is desirable to use protein concentrations not greater than about 10 times the *K*_d_ value.^[34]^ The brightness of 4-CN-Trp fluorescence thus offers the important advantage that it is sufficiently bright for facile detection at nanomolar protein concentrations, which also helps alleviating the inner-filter effect associated with the absorbance of free ligand. The similarity in dissociation constant observed for FKBP12 and FKBP12-4CNW is another example of how little the characteristics of a protein may change when a Trp residue is changed to a 4-CN-Trp residue.

It has been reported that the wavelength of the fluorescence emission maximum of 4-CN-Trp depends on the polarity of the environment, varying between 395 nm in THF and 425 nm in water,^[9]^ and the wavelength of the emission maximum in FKBP12-4CNW also differs between the free protein and the complex with rapamycin (Figure S19b). Provided the line shapes of both states can be fitted reliably, decomposition of the fluorescence signal into its individual components may present an alternative way of measuring *K*_d_ values, which is less sensitive to inner filter effects and collision quenching.

### Determining the Site of Ligand Binding

The quenching of tryptophan fluorescence by aromatic ligands has been used to determine, which Trp residue in a protein is most prominently involved in the interaction with the ligand and therefore nearest to the ligand binding site.^[35]^ As the fluorescence spectra of the individual Trp residues are not resolved, the experiment requires eliminating their contributions by site-specific mutagenesis. However, even substituting tryptophan by other aromatic amino acids, such as phenylalanine or tyrosine, presents a significant alteration that risks affecting the protein structure and ligand binding. Furthermore, removing the contribution of a single tryptophan residue changes the overall fluorescence profile relatively little if the protein contains multiple tryptophan residues. A much better approach is to replace a single Trp residue by 4-CN-Trp, which minimally alters the protein structure and the fluorescence of which can be selectively excited without interference from any of the remaining Trp residues.

To illustrate the approach, we use the ligand binding domain (LBD) of the human nuclear xenobiotic pregnane X receptor (hPXR) which promiscuously binds a large number of synthetic drugs in a deep ligand binding pocket.^[36]^ The wild-type protein of the LBD contains three Trp residues, where Trp199 and Trp223 are highly solvent-exposed, and Trp299 lines the ligand binding pocket. We produced three mutants of the LBD, replacing the Trp residues one by one by a 4-CN-Trp residue. As expected, the emission wavelengths of the solvent-exposed CN-Trp residues were longer than for the partially buried CN-Trp residue in position 299 (Figure 6a).

**Figure 6.**
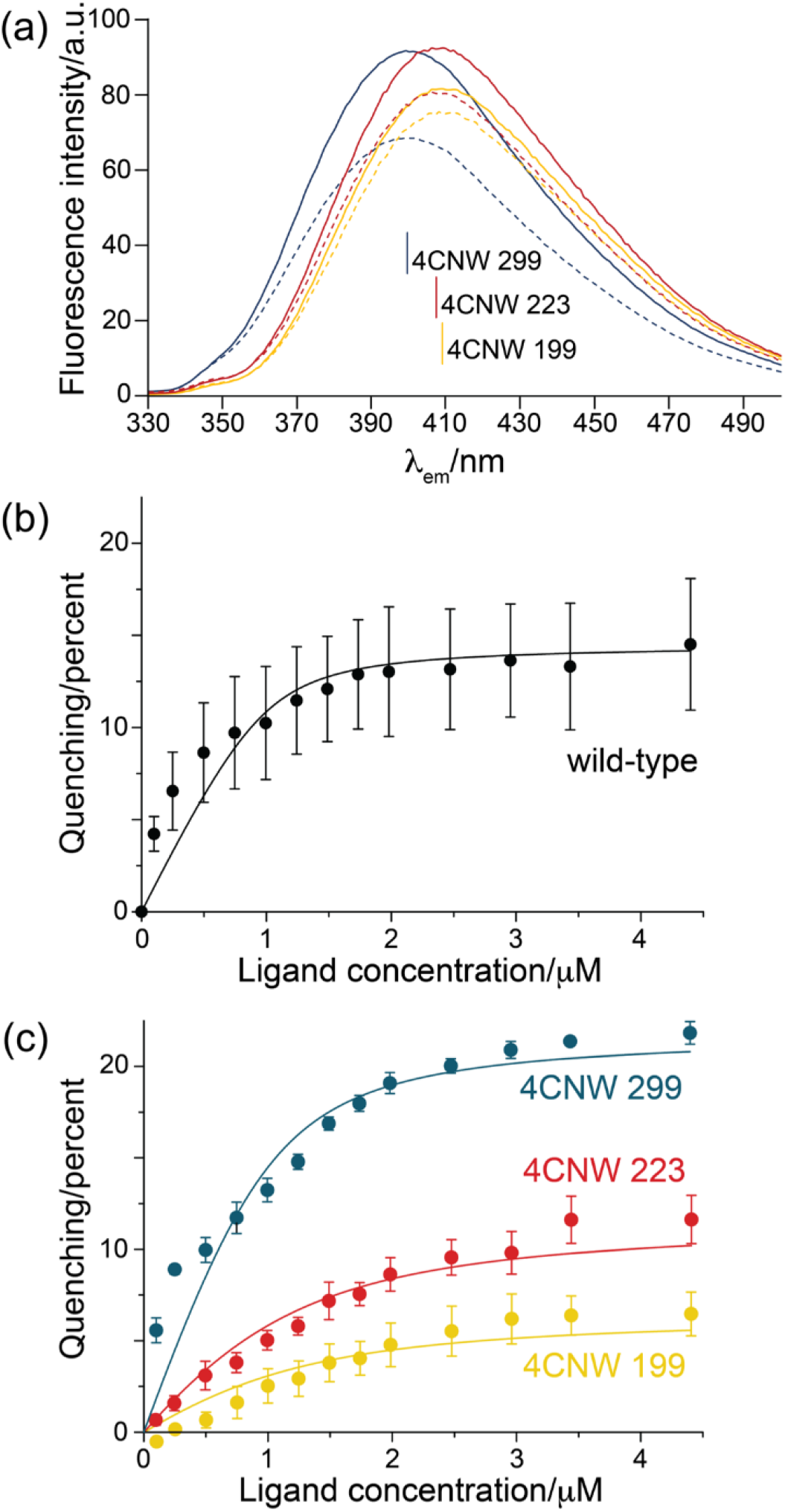
Fluorescence quenching experiments of the hPXR-LBD with **1**. (a) Fluorescence emission spectra (excitation wavelength 310 nm) of 4-CN-Trp mutants in positions 199 (dark blue), 223 (red), and 299 (yellow). The spectra were recorded at room temperature of 1 μM solutions without (solid lines) and with (dashed lines) 5.5 μM **1**. The buffer contained 50 mM Tris-HCl, pH 7.5, and 300 mM NaCl. (b) Fluorescence quenching of Trp observed in an experiment titrating the hPXR LBD with **1**. Fluorescence intensity was recorded with excitation at 280 nm and detection at 335 nm. The inner filter effect was accounted for by subtracting the spectra obtained with **1** only. The fitted two-state binding curve corresponds to *K*_d_ = 90 nM. Error bars are standard deviations and were determined from performing the experiment in triplicate. (c) Same as (b), except that the fluorescence was excited at 310 nm and detected at the emission maxima of the 4-CN-Trp mutants (see (a)). The fits correspond to *K*_d_ values of 600 nM at site 199, 500 nM at site 223, and 170 nM at site 299. The error bars are smaller than for the Trp fluorescence in (b) because the greater brightness of 4-CN-Trp allowed setting a smaller gain on the fluorimeter.

Using TO901317 (compound **1**, Figure S20)^[37]^ as the ligand in a titration experiment with the wild-type protein resulted in quenching of the Trp fluorescence (excitation at 280 nm, detection at 335 nm) with increasing amounts of **1**, allowing an estimate of the dissociation constant (Figure 6b). The co-crystal structure of hPXR-LBD with **1** (PDB ID 2O9I)^[38]^ shows that the shortest distances between the aromatic rings in the ligand and Trp199, Trp223, and Trp299 are 16 Å, 12 Å, and 3 Å, respectively, suggesting that Trp199 contributes the least and Trp299 the most to the fluorescence quenching effect. To test this prediction, we titrated the hPXR-LBD mutants with **1** and monitored the fluorescence emission at 410 nm with steady-state excitation at 310 nm. As expected, the mutant with 4-CN-Trp in position 299 revealed the largest fluorescence quenching effect, whereas the mutants with 4-CN-Trp in positions 199 and 223 showed effects of smaller magnitude, which were less readily fitted by the equation for two-state binding (Figure 6c). This result clearly demonstrates the closer proximity of the ligand to Trp299 than the other Trp residues. The smaller but still significant fluorescence quenching effects observed for the mutants in positions 199 and 223 may be governed by collision-induced quenching by free ligand,^[4]^ as these residues are highly solvent exposed. Also in position 299, the fluorescence quenching effect may be enhanced by ligand-induced protection from hydration water.

In the case of TO901317 binding to hPXR, a previously used radioactive assay with ^3^H-labeled **1** binding to hPXR tethered to beads yielded *K*_d_ = 26 ±3 nM,^[37]^ while subsequent inhibition experiments of the hPXR-LBD fused to GST determined IC50 values ranging from 52 to 159 nM using a time-resolved FRET assay.^[39]^ The *K*_d_ value determined in our work by straightforward titration of hPXR-LBD with **1** (170 nM) thus suggests that the presence of the cyano group does not disrupt the binding mode, although the binding affinity is reduced. Notably, however, the Trp site contributing most strongly to the fluorescence quenching effect can be identified already from a semi-quantitative evaluation of the titration data.

## Conclusion

The fluorescence applications demonstrated in the present work establishes 4-CN-Trp as an exceptionally useful probe for exploring ligand binding. First, the sensitivity of the fluorescence of 4-CN-Trp can be used to measure sub-nanomolar ligand binding affinities. These measurements can be performed conveniently in isotropic solution without immobilizing the protein or target on a solid support. Second, the quenching of the fluorescence of 4-CN-Trp by aromatic ligands can be used to determine their binding site on target proteins containing several Trp residues. Importantly, the fluorophore can be buried inside the protein without significantly altering the protein structure. Our results align with the finding by Talukder et al., who reported functional conservation of enzymatic activity for dihydrofolate reductase labeled with 6-CN-Trp and 7-CN-Trp.^[13]^

The aminoacyl-tRNA synthetases of the present work enable site-selective installation of 4-CN-Trp and three more CN-Trp isomers in proteins in response to an amber stop codon with excellent specificity and yield, delivering pure proteins without misincorporation of canonical amino acids at the amber stop codon or premature chain termination.

Combined with the published protocol for the enzymatic conversion of CN-indoles into the corresponding CN-Trp isomers,^[24]^ the high-yield in vivo protein production with any of these CN-Trp variants has become affordable and readily achievable in standard biochemical laboratories.

With these advances, the spectroscopic applications of CN-Trp isomers developed over the past few decades and demonstrated with synthetic peptides are now effortlessly applicable to proteins, opening the door to many exciting applications. Among these, the potential uses of 4-CN-Trp are particularly promising, ranging from measurements of local electrostatic fields and H-bonds,^[40]^ FRET and photo-induced electron transfer between 4-CN-Trp and tryptophan for proximity measurements^[41]^ to two-photon microscopy,^[42]^ and straightforward microscopy, e.g., for the intracellular tracking of polypeptides.^[11, 17]^ Our present results show that 4-CN-Trp fluorescence can readily be excited using the 355 nm excitation laser commonly found in fluorescence microscopy and flow cytometry instruments.

To encourage the uptake of this technology, the requisite plasmids have been deposited at Addgene (Watertown, MA) (#222871, #222872, #222873, and #222874).

## Supporting Information

The authors have cited additional references within the Supporting Information.^[43–55]^

## Acknowledgements

We thank Dr. Harpreet Vohra and Michael Devoy at the John Curtin School of Medical Research, Australian National University for technical support on the FACS experiments. We appreciate Prof. Shigeki Kiyonaka at Nagoya University for his constructive advice on FKBP12-related experiments. We acknowledge Microscopy Australia (ROR: 042mm0k03) at the Centre for Advanced Microscopy, Australian National University, especially, Daryl Webb, for his knowledge and input. This research was undertaken in part using the MX2 beamline at the Australian Synchrotron, part of ANSTO, and made use of the Australian Cancer Research Foundation (ACRF) detector. Financial support by the Australian Research Council, including the Centre of Excellence for Innovations in Peptide and Protein Science (CE200100012) and Discovery Projects (DP230100079, DP210100088) is gratefully acknowledged.

