## Supporting Information for "Rendering Proteins Fluorescent Inconspicuously: Genetically Encoded 4-Cyanotryptophan Conserves Their Structure and Enables the Detection of Ligand Binding Sites"

#### Table of contents:

**Methods**

a) Materials
b) Selection of functional G1PylRS enzymes recognizing cyanotryptophans
c) Substrate polyspecificity analysis of selected G1PylRS mutants
d) In vivo protein expression and purification of FKBP12, hPXR-LBD, and MBP
e) ESI mass spectrometry intact protein analysis
f) Live cell imaging
g) Fluorescence measurements
h) Ligand titration experiments
i) Protein crystallization and structure determination

**Supplementary figures**

**Figure S1–S20**

**Supplementary tables**

**Table S1–S4**

**References**

### Methods

#### a) Materials

No unexpected or unusually high safety hazards were encountered. The non-canonical amino acids (ncAA) 4-cyano-L-tryptophan (4-CN-Trp, Cat. ID: V152071), 5-cyano-L-tryptophan (5-CN-Trp, Cat. ID: M22603), 6-cyano-L-tryptophan (6-CN-Trp, Cat. ID: V150864), and 4-bromo-L-tryptophan (4-Br-Trp, Cat. ID: M22628) were purchased from Advanced ChemBlocks Inc. (USA). 7-Cyanoindole (Cat. No.: A378351), 4-fluoro-DL-tryptophan (4-F-Trp, Cat. No.: A901033), 4-chloro-L-tryptophan (4-Cl-Trp, Cat. No.: A710963), 4-methyl-DL-tryptophan (4-Me-Trp, Cat. No.: A170319), 5-fluoro-DL-tryptophan (5-F-Trp, Cat. No.: A114530), 5-chloro-L-tryptophan (5-Cl-Trp, Cat. No.: A146648), 5-bromo-L-tryptophan (5-Br-Trp, Cat. No.: A428010), 5-methyl-DL-tryptophan (5-Me-Trp, Cat. No.: A767222), 5-hydroxy-L-tryptophan (5-HTP, Cat. No.: A102374), 5-methoxy-L-tryptophan (5-MTP, Cat. No.: A228826), 6-fluoro-DL-tryptophan (6-F-Trp, Cat. No.: A339425), 6-chloro-L-tryptophan (6-Cl-Trp, Cat. No.: A124813), 6-bromo-L-tryptophan (6-Br-Trp, Cat. No.: A499754), 6-methyl-L-tryptophan (6-Me-Trp, Cat. No.: A521841), 7-fluoro-DL-tryptophan (7-F-Trp, Cat. No.: A158059), 7-methyl-DL-tryptophan (7-Me-Trp, Cat. No.: A743946), and rapamycin (Cat. No.: A656002) were purchased from Ambeed (USA). TO901317 (Cat. No.: T2320) was purchased from Merck (Germany).

#### b) Selection of functional G1PylRS enzymes recognizing cyanotryptophans

The selection used a previously established library of G1PylRS mutants on the plasmid pBK-G1RS transformed into *E. coli* DH10B cells harboring the reporter plasmid pBAD-H6RFP. [19a] Following recovery from transformation, the culture was directly inoculated into a flask with 25 mL LB medium containing 100 mg/L carbenicillin and 50 mg/L kanamycin, supplied with 0.4% L-arabinose and 1 mM ncAA, which served as the sample for the first round of positive selection (**1P+**). Overnight expression at 37 °C led to a readily detectable level of RFP expression. Cells were resuspended in 5 mL PBS buffer (137 mM NaCl, 2.7 mM KCl, 10 mM $Na_2HPO_4$, 1.8 mM $KH_2PO_4$, pH 7.4) after harvesting. A 100-fold dilution yielded a concentration suitable for fluorescence-activated cell sorting (FACS) on an FACSAria Fusion cell sorter (BD Biosciences, USA; Figure S4).

Cells with high RFP levels were collected from the **1P+** sample (1.0% for 4-CN-Trp, 0.3% for 5-CN-Trp, 0.4% for 6-CN-Trp, and 0.7% for 7-CN-Trp as indicated by violet shades in Figure S4) and subjected to a following round of negative selection. Without the addition of ncAAs, the cells were regrown as sample **2N–**, from where cells with low RFP expression levels (75.0% for 4-CN-Trp, 52.4% for 5-CN-Trp, 58.8% for 6-CN-Trp, and 44.4% for 7-CN-Trp) were collected. These cells were aliquoted to inoculate media with positive (**3P+**) and negative (**3P–**) conditions. The cell population with high RFP fluorescence in the **3P**+ sample in the selection attempt for 4-CN-Trp was obviously greater than in the **3P**– sample, indicating the successful accumulation of active G1PylRS variants specific for 4-CN-Trp. The top 3.3% RFP fluorescent cells of the **3P+** sample for 4-CN-Trp were collected for final analysis. The top RFP cells from the other **3P+** samples were collected (9.4% for 5-CN-Trp, 3.8% for 6-CN-Trp, and 4.7% for 7-CN-Trp) and cultured for the fourth round without cyanotryptophans. The cells showing low RFP expression levels in the **4N**– samples (48.5% for 5-CN-Trp, 29.4% for 6-CN-Trp, and 53.1% for 7-CN-Trp) were collected and again aliquoted to inoculate media with positive (**5P+**) and negative (**5P–**) conditions. These cultures showed a clear response to the presence of the ncAAs and the top 3.3%–3.4% RFP fluorescent cells were collected from the **5P+** samples. From each of the 4-CN-Trp, 5-CN-Trp, 6-CN-Trp, and 7-CN-Trp selection experiments, an aliquot of 2,000 cells were allowed to recover on LB agar plates containing 100 mg/L carbenicillin and 50 mg/L kanamycin, and isolated colonies were analyzed using 96-well plates. Cells containing 60 enzyme candidates were inoculated into both positive (with 1 mM CN-Trp) and negative (without ncAA) growth conditions. The intensity of red fluorescence was measured as an indicator of the expression level of amber-interrupted RFP after expression overnight, using a TECAN Infinite 200 Pro M Plex plate reader (Tecan, Switzerland) and normalized by the $OD_{600}$ value of the cell culture. Sequences of four candidates from the 4-CN-Trp selection, three from the 5-CN-Trp selection, two from the 6-CN-Trp selection, and three from the 7-CN-Trp selection were identified as novel G1PylRS mutants incorporating the respective cyanotryptophans. Table S2 lists the amino acid mutation sets found.

**c) Substrate polyspecificity analysis of selected G1PylRS mutants**

The genes of the four selected G1PylRS mutants 4CNWRS, 5CNWRS, 6CNWRS, and 7CNWRS were subcloned into the high-copy number plasmid pRSF plasmid. The plasmid map of pRSF-4CNWRS is shown in Figure S8. The resulting pRSF plasmids were individually co-transformed into *E. coli* B-95.ΔAΔ*fabR* cells[28] with the reporter plasmid pCDF-H6RFP encoding a $His_6$-TAG-RFP gene under the control of a T7 promoter. Cells were grown in 10 mL LB medium containing 50 mg/L spectinomycin and 50 mg/L kanamycin at 37 $^oC$. 20 μL of each overnight culture were used to inoculate 160 μL LB medium grown on 96-well plates, supplemented with the same antibiotic, along with 1 mM IPTG under different conditions of ncAA supplementation. Each culture was prepared in triplicate. The plates were shaken at 400 rpm for 16 h at 30 $^oC$ and kept at 4 $^oC$ for 12 h for complete expression and maturation of RFP. The intensities of red fluorescence measured on the plate reader were normalized by the $OD_{600}$ value of the cell culture and the average of triplicates was determined.

**d) In vivo protein expression and purification of FKBP12, hPXR-LBD, and MBP**

pCDF plasmids containing the genes of wild-type and amber-interrupted FKBP12, hPXR LBD, and MBP were purchased from Twist Bioscience (USA). The FKBP12 and hPXR LBD mutants contained a C-terminal $His_6$ tag. The MBP mutants contained a N-terminal $His_6$ tag followed by a tobacco etch virus (TEV) protease recognition site. The plasmid map of pCDF-FKBP12-W59TAG is shown in Figure S9.

For the site-specific incorporation of 4-CN-Trp, proteins were produced in *E. coli* B-95.ΔAΔ*fabR* cells[28] co-transformed with pRSF-4CNWRS and the pCDF plasmid containing the amber codon-interrupted gene of the respective target protein. The transformed cells were grown overnight at 37 °C in LB medium containing 25 mg/L kanamycin and 25 mg/L spectinomycin. Overnight cultures were inoculated into fresh LB medium (1:100 dilution), supplemented with 25 mg/L kanamycin, 25 mg/L spectinomycin, 1 mM 4-CN-Trp. After the cultures reached an $OD_{600}$ value of 0.6–1, protein expression was induced by the addition of 1 mM isopropyl-β-D-thiogalactopyranoside (IPTG). The cells were harvested by centrifugation following overnight expression at room temperature. Wild-type hPXR-LBD and MBP were produced in the same way, following transformation with the pCDF vector containing the gene of wild-type hPXR-LBD or MBP and bacterial growth in the absence of kanamycin and 4-CN-Trp.

To purify the proteins, the cell pellet was resuspended in buffer A (50 mM Tris-HCl pH 7.5, 300 mM NaCl, 5% glycerol, 10 mM imidazole) and lysed using an Avestin Emulsiflex C5 system (Avestin, Canada) using two passes with a pressure of 10,000–15,000 psi. The lysate was clarified by centrifugation (30,000 *g*, 60 min, 4 °C) and loaded onto a 1 mL His GraviTrap column (Cytiva, USA). The column was washed with 20 column volumes buffer B (same as buffer A but with 20 mM imidazole) and the protein was eluted with 5 column volumes buffer C (same as buffer A but with 500 mM imidazole). The eluted proteins were dialyzed against buffer D (same as buffer A but without imidazole and glycerol) at 4 °C and concentrated using an Amicon ultracentrifugation centrifugal tube with a molecular weight cutoff (MWCO) of 10 kDa. For FKBP12-W59-4CNW sample, protein was dialyzed against PBS buffer. For the MBP samples, protein was dialyzed against buffer E (50 mM Tris-HCl pH 7.5, 300 mM sodium chloride, 1 mM dithiothreitol (DTT)), and TEV proteolytic cleavage was done by incubation with TEV protease at 4 °C for 16 h. The cleaved MBP samples were recovered by a reverse IMAC step and dialyzed against buffer D. Yields of hPXR-LBD mutants produced with 4-CN-Trp ranged between 7 and 10 mg of purified protein per liter of cell culture. For comparison, the yield of wild-type hPXR-LBD was 30 mg per liter of cell culture. Yields of MBP mutants with 4-CN-Trp ranged between 10 and 20 mg of purified protein per liter of cell culture. The yield of FKBP12 with 4-CN-Trp was 63 mg of purified protein per liter of cell culture.

**e) ESI mass spectrometry intact protein analysis**

Intact protein mass spectrometry was performed on an Orbitrap Elite Hybrid Ion Trap-Orbitrap mass spectrometer (Thermo Fisher Scientific, USA) connected to a Thermo Fisher Scientific UltiMate 3000 HPLC system equipped with ZORBAX 300SB-C3, 3.5 µm, 4.6 x 50 mm HPLC column (Agilent Technologies, USA). Approximately 7.5 pmol of sample was injected using a 500 µL/min linear gradient of solvent A (0.1% (v/v) formic acid in water) and solvent B (0.1% (v/v) formic acid in acetonitrile), ramping solvent B from 5% solvent B at the start to 80% after 12 min. Data were collected using an electrospray ionization (ESI) source in positive ion mode. Protein intact mass was determined by deconvolution using the program Xcalibur 3.0.63 (Thermo Fisher Scientific, USA).

**f) Live cell imaging**

For fluorescence microscopy, *E. coli* B-95.ΔAΔ*fabR* cells were co-transformed with pRSF-4CNWRS and the pCDF plasmid containing one of the amber codon-interrupted genes FKBP12-W59TAG, Intimin-TAG, OmpX-TAG, or PelB-NB-F67TAG (see Table S3 for the nucleotide and amino acid sequences). To check the baseline level of free 4-CN-Trp non-covalently associating with the cells, a control sample was prepared using pRSF and pCDF plasmids lacking the genetic encoding system for 4-CN-Trp. The cells were cultured in 50 mL LB medium with 25 mg/L kanamycin and 25 mg/L spectinomycin at 37 °C until reaching an $OD_{600}$ of 0.4. Each culture was then split into two flasks and one of them supplemented with 0.75 mM 4-CN-Trp. Protein expression was induced with 1 mM IPTG and the cells were incubated overnight at room temperature.

To remove free 4-CN-Trp associated with the cells, the cells were washed three times with PBS buffer and subsequently incubated on ice for 12 hours. Ten microliters of the resuspended cells (diluted three-fold compared to the overnight culture) were loaded onto 1% agarose pads and covered with coverslips. Samples were imaged using a Zeiss LSM780 UV-NLO confocal microscope (Carl Zeiss Microscopy GmbH, Germany) with a 355 nm laser for excitation. Emission signals below 503 nm were recorded with the same gain and laser power for all samples. The focus was adjusted based on the fluorescence channel and the transmission channel was recorded simultaneously to capture the brightfield image. Image processing was performed using ZEN lite 3.10 software (Carl Zeiss Microscopy GmbH, Germany). Statistical analysis of the imaged cells was conducted on a FACSAria Fusion cell sorter (BD Biosciences, USA) using a 355 nm laser and a 379/28 bandpass filter.

**g) Fluorescence measurements**

The fluorescence data of Figure 3/4 were obtained using a CARY Eclipse fluorescence spectrophotometer (Agilent, USA) with a 3 mL/400 μL quartz cuvette. The excitation and emission slit widths were set to 5 nm and 5 nm (Figure 3a), 10 nm and 20 nm (Figure 3c), or 5 nm and 10 nm (Figure 4), respectively. PBS buffer (or buffer E as required) was used as the blank. The fluorescence of Trp and 4-CN-Trp was measured with excitation wavelengths of 280 nm and 310 nm, respectively. For measurements at low protein concentrations, it proved critical to thoroughly clean the quartz cuvette prior to the measurements. This was achieved by incubating the cuvette with 2% detergent Hellmanex™ in Milli-Q water at 45 °C for 20 min and repeating this process until refilling of the cuvette with PBS buffer gave reproducible background intensity. The data points of Figure 3b were obtained by scanning the emission spectra with excitation at 310 nm using the photomultiplier tube (PMT) detector voltage at 950 V. The excitation and emission slit were 10 nm and 20 nm respectively. The scan rate was 600 nm/min with 1 nm of data interval. For each emission spectrum, two scans were taken and averaged to minimize fluctuations in the detection of fluorescence intensity. Following subtraction of the background signal measured without protein, the signal was smoothed by averaging five data points, up to 2 nm on either side of 380 nm and spaced by 1 nm. For Figure 4, the signal-to-noise ratio was improved in the same way by averaging between ten measurements at 380 nm and up to 2 nm on either side.

**h) Ligand titration experiments**

The titration data of Figure 3c were obtained in the CARY fluorimeter by titrating 500 nM – 10 μM stock solutions of rapamycin into a 4 nM solution of FKBP12-4CNW in PBS buffer at room temperature. The solution was allowed to equilibrate for 6 minutes before reading the fluorescence. The titration was repeated three times and each data point representing the average of two measurements. A single set of two measurements, recorded with the same settings, was taken for the background measurements at the corresponding concentrations of rapamycin in PBS buffer, which were then used for baseline subtraction.

The titration data of Figure 4 were obtained in the CARY fluorimeter by titrating 0.1 mM – 0.25 mM stock solutions of **1** into a 1 μM solution of the hPXR-LBD in buffer E at room temperature. The solution was allowed to equilibrate for 15 minutes before reading the fluorescence. Data from three independent experiments were analyzed. To correct for the inner filter effect, the titration was repeated with 1 μM solutions of Trp or 4-CN-Trp in buffer E.

$K_d$ values were determined by fitting the titration data using the equation

$$\Delta IF = \frac{(K+Pt+Lt) - \sqrt{(K+Pt+Lt)^2 - 4Lt\,Pt}}{2\,Pt} \qquad (1)$$

where $\Delta IF$ is the change in fluorescence intensity, $K$ the equilibrium dissociation constant, and $Pt$ and $Lt$ denote the total protein and ligand concentrations, respectively.

**i) Protein crystallization and structure determination**

The wild-type MBP (MBP-WT) sample was loaded onto a HiLoad 26/600 Superdex 75 pg column (Cytiva, USA) equilibrated with buffer F (10 mM MES pH 6.2, 0.2% sodium azide). The fractions containing MBP-WT were identified using SDS-PAGE and combined and concentrated to 35 mg/mL using a 10 kDa MWCO Amicon ultrafiltration centrifugal tube (Merck Millipore, USA). The target protein was diluted to 15 mg/mL in buffer F. Crystallization condition screening was performed using a sparse matrix screen Crystal Screen HT (Hampton Research) at 18 ˚C with 1:1 ratio of protein:reservoir solution in a total volume of 0.5 µL. Multiple crystals of sufficient size grew in a solution containing 0.1 M cadmium chloride, 30% PEG 400, and 0.1 M sodium acetate pH 4.6 (condition E12) within 36 hours. Crystals of MBP-WT were reproduced using hanging drop vapor diffusion of 1 µL protein and 2 µL well solution against a 500 µL reservoir of E12 buffer.

After the reverse IMAC step, the MBP mutant with 4-CN-Trp (MBP-4CNW) was dialyzed into buffer F and concentrated to 15 mg/mL using a 10 kDa MWCO Amicon ultrafiltration centrifugal tube (Merck Millipore, USA). MBP-4CNW crystals were obtained by seeding crystals of the MBP-WT protein. 3 µL of the MBP-WT crystal drop was pipetted into an Eppendorf tube containing 50 µL of condition E12 and crushed using a seed crusher made from a melted pipette tip. This solution was then diluted to 550 µL to produce crystal seeding well solution. Serial dilutions were made to test for optimal seed concentration. Large, cubic protein crystals were obtained within 36 hours by equilibrating drops of 1 µL MBP-4CNW sample and 2 µL well solution against a reservoir containing $10^6$ dilution of seed stock in 30% PEG 400, and 0.1 M sodium acetate pH 4.6. A single crystal was mounted and cryoprotected using Parabar (Hampton Research) before flash-freezing in liquid nitrogen.

For both MBP-4CNW and MBP-WT crystals, diffraction data were collected at 100 K at the MX2 beamline at the Australian Synchrotron,[43] and processed using XDS in the P 1 $2_1$ 1 space group.[44] Data were scaled in Aimless (CCP4)[45] until the data was of sufficient quality in the highest resolution shell. The structure was solved using molecular replacement (Phaser MR,[46] CCP4) using PDB ID 1JW4 as the search model, followed by remodeling in Coot,[47] and refinement in Phenix.[48] For both crystal structures, the space group was validated using Zanuda. [49] The identity and the coordination environment of the Na and Cd ions was validated using the CheckMyMetal server.[50]

For the MBP-4CNW structure, CIF restraints for the non-canonical amino acid were generated by adding the cyano-group to a Trp amino acid in PyMOL[51] and generating hydrogens, followed by processing of the resulting PDB file in eLBOW.[52] The resulting CIF dictionary was then modelled in the structure using the replace residue function in Coot, and used as a restraint for refinement.

Data collection and refinement statistics are given in Table S4. The completed structures are available in the Protein Data Bank with accession IDs 9CLD (MBP-WT) and 9CLC (MBP-4CNW).

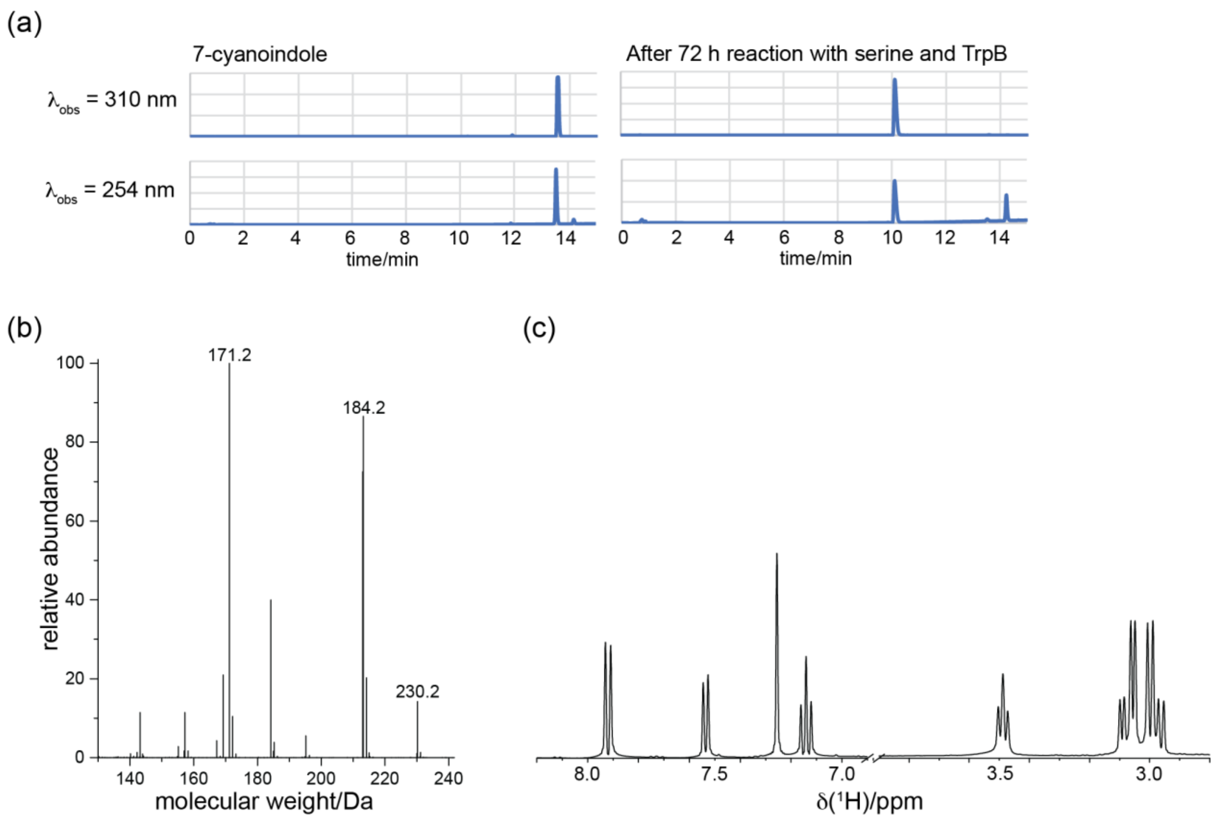

**Figure S1.** Analytical data demonstrating the purity of 7-cyanotryptophan produced by incubation of serine with 7-cyanoindole and the TrpB mutant *Tm*9D8*.[24] (a) Left panel: HPLC traces of 7-cyanoindole. Right panel: after 72 h reaction with serine and the TrpB mutant at 55 °C and spontaneous precipitation of 7-CN-Trp after cooling of the reaction mixture. HPLC analysis was performed on a C-18 silica column (2.1 × 100 mm 1.8-Micron) using a gradient of acetonitrile/water (each containing 0.1% formic acid by volume) starting from 5% to 90% acetonitrile over 15 min at 0.5 mL/min. An NMR spectrum of the reaction mixture recorded at the end of the reaction showed no evidence of any remaining 7-CN-indole. (b) Mass spectrum of precipitated 7-CN-Trp (calculated $[M+H]^+$ = 230.2). (c) $^1$H NMR spectrum of 1 mM solution of the precipitant recorded in PBS + 10% $D_2O$ at 400 MHz. δ/ppm: 7.92 (H4), 7.55 (H6), 7.25 (H2), 7.14 (H5), 3.49 ($H^{\alpha}$), 3.08 and 2.98 ($C^{\beta}H_2$).

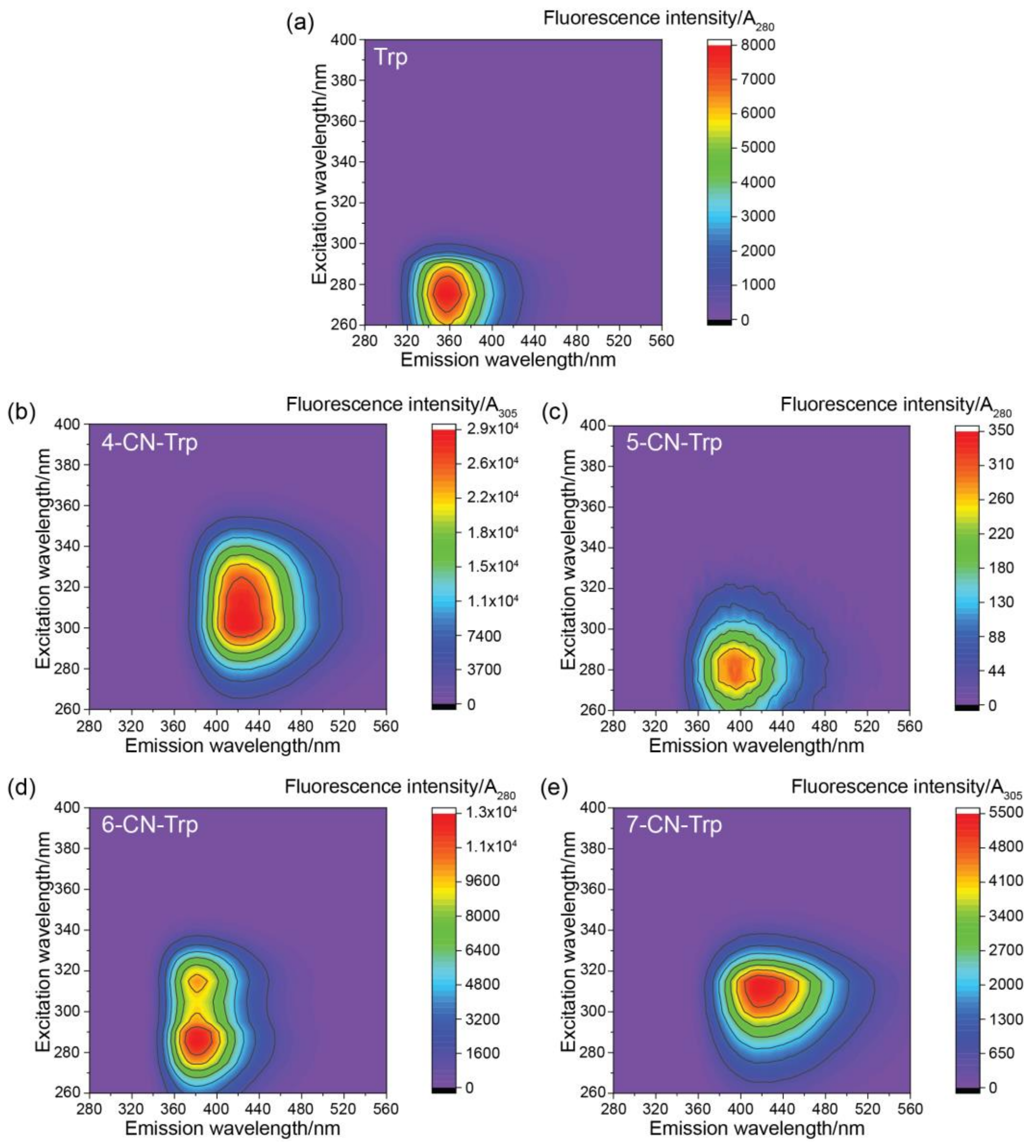

**Figure S2.** Fluorescence spectra of aqueous solutions of tryptophan and 4-, 5-, 6-, and 7-cyanotryptophan in PBS solution at 25 °C. The absorption spectra were measured using a Cary 60 UV-Vis spectrophotometer (Agilent, USA). The fluorescence spectra were measured using a Cary Eclipse Fluorescence spectrophotometer (Agilent, USA) with 5 nm excitation and emission slits and PMT voltage of 600 V. To record the fluorescence with the same gain setting, the tryptophan compounds were used at different concentrations (16 μM Trp, 3 μM 4-CN-Trp, 16 μM 5-CN-Trp, 10 μM 6-CN-Trp, 10 μM 7-CN-Trp) while keeping the UV absorbance below 0.1 at the excitation wavelength of each sample to minimize the self-quenching effect. The emission signal intensities are normalized by the intensity of the absorption maximum (280 nm for Trp, 5-CN-Trp, and 6-CN-Trp; 305 nm for 4-CN-Trp and 7-CN-Trp). (a) Tryptophan. (b) 4-CN-Trp. (c) 5-CN-Trp. (d) 6-CN-Trp. (e) 7-CN-Trp.

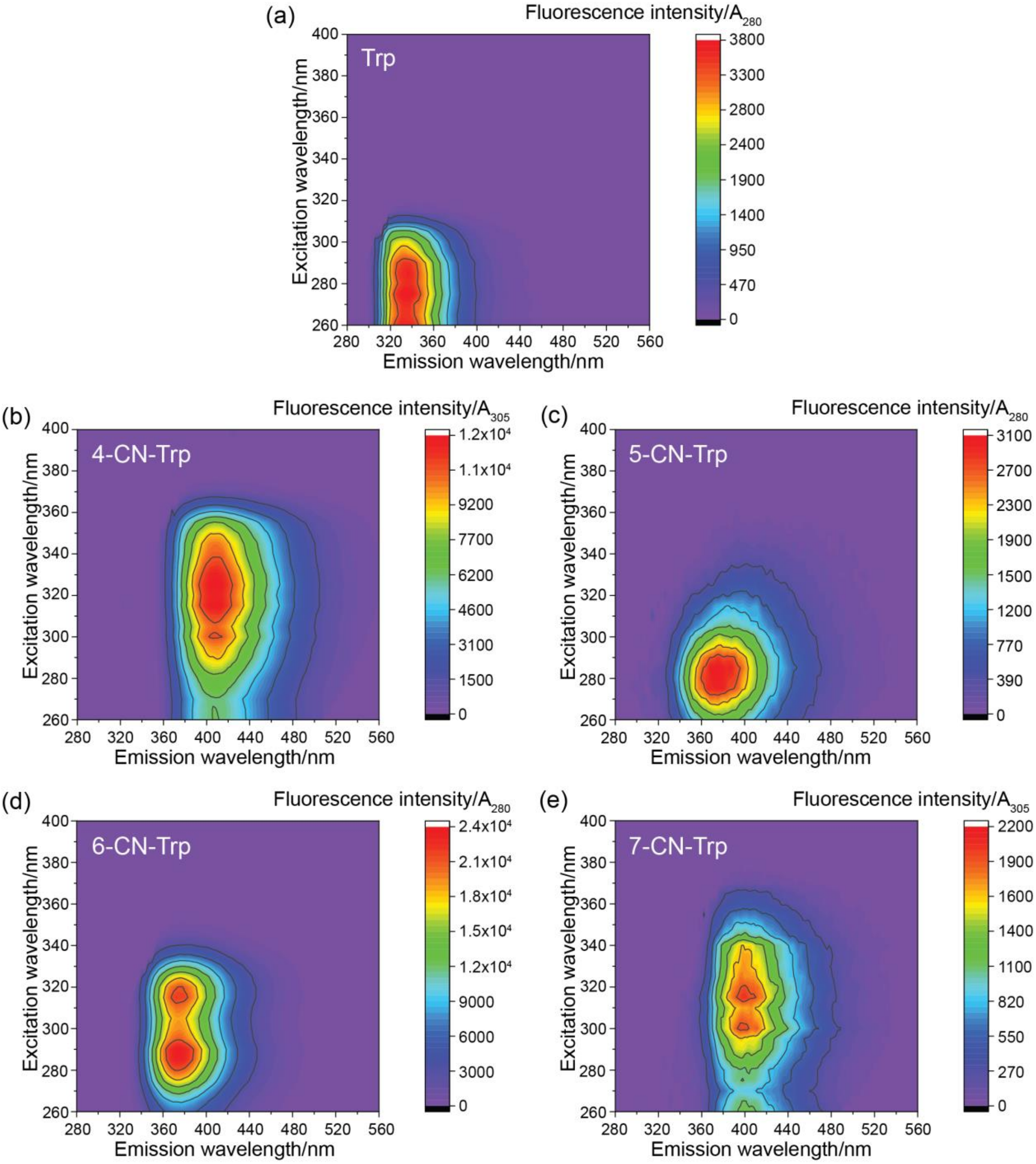

**Figure S3.** Fluorescence spectra of tryptophan and 4-, 5-, 6-, and 7-CN-Trp solutions (same concentration as in Figure S2) in 100% acetonitrile at 25 ºC. Except for the solvent, all parameters were as in Figure S2. (a) Tryptophan. (b) 4-Cyanotryptophan. (c) 5-Cyanotryptophan. (d) 6-Cyanotryptophan. (e) 7-Cyanotryptophan. The distinct fluorescence characteristics of the CN-Trp isomers are also reflected in the corresponding cyanoindoles, for which much more detailed photophysical characterizations have been carried out.[11, 53, 54]

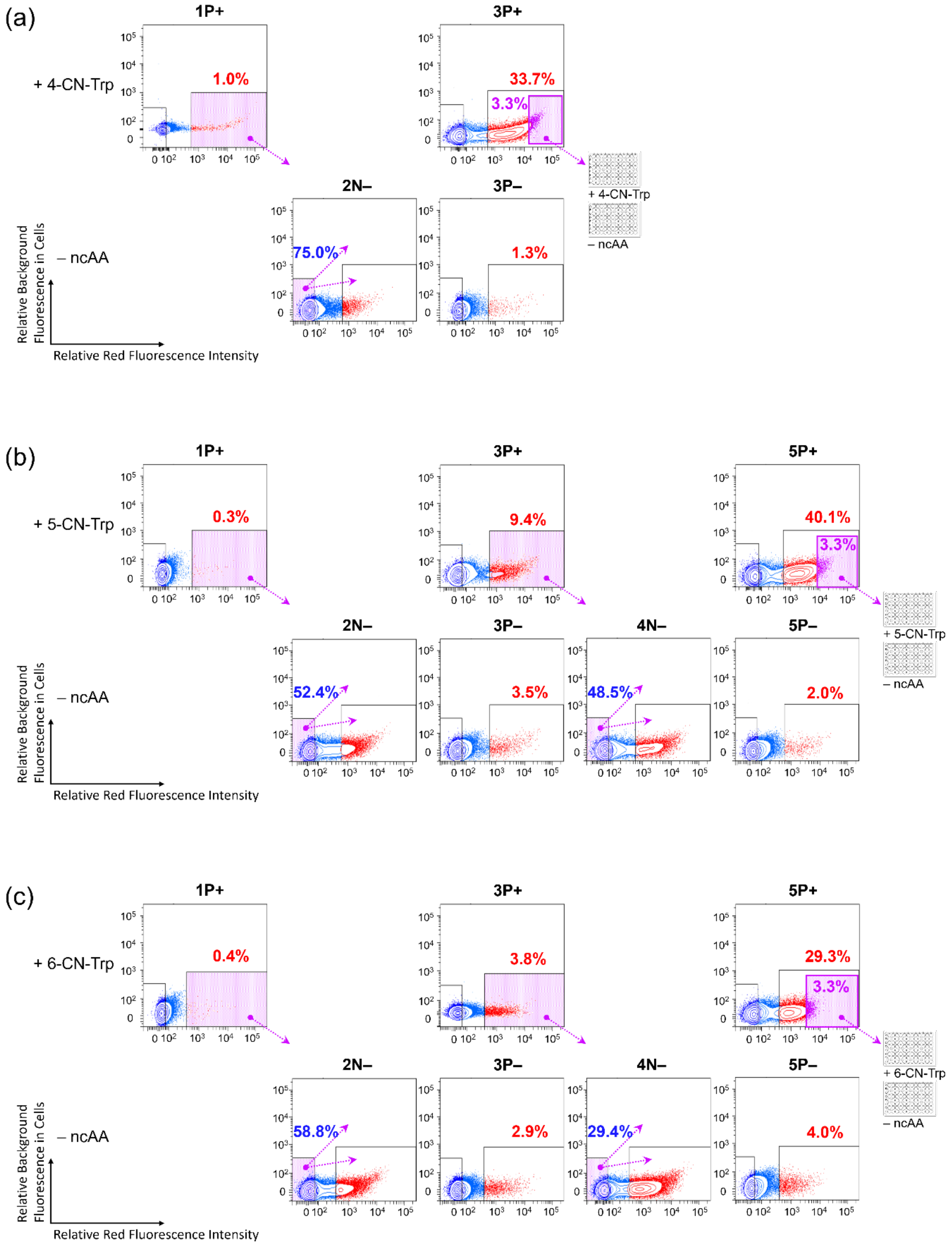
(a)
1P+
3P+
+ 4-CN-Trp
1.0%
33.7%
3.3%
2N–
3P–
– ncAA
75.0%
1.3%
+ 4-CN-Trp
– ncAA
Relative Background Fluorescence in Cells
Relative Red Fluorescence Intensity
(b)
1P+
3P+
5P+
+ 5-CN-Trp
0.3%
9.4%
40.1%
3.3%
2N–
3P–
4N–
5P–
– ncAA
52.4%
3.5%
48.5%
2.0%
+ 5-CN-Trp
– ncAA
Relative Background Fluorescence in Cells
Relative Red Fluorescence Intensity
(c)
1P+
3P+
5P+
+ 6-CN-Trp
0.4%
3.8%
29.3%
3.3%
2N–
3P–
4N–
5P–
– ncAA
58.8%
2.9%
29.4%
4.0%
+ 6-CN-Trp
– ncAA
Relative Background Fluorescence in Cells
Relative Red Fluorescence Intensity

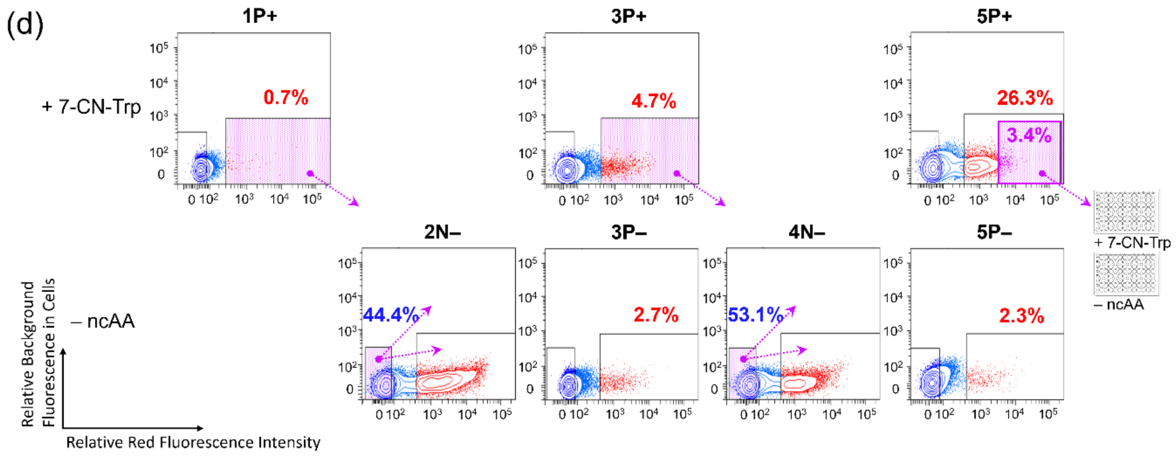

**Figure S4.** Results from fluorescence-activated cell sorting (FACS) screening for active and specific G1PylRS enzymes that recognize (a) 4-CN-Trp, (b) 5-CN-Trp, (c) 6-CN-Trp, or (d) 7-CN-Trp. The horizontal axis indicates the intensity of red fluorescence following excitation at 560 nm. The vertical axis plots the level of background fluorescence in cells excited at 488 nm. **P** and **N** identify positive and negative selection rounds, respectively. + and – labels identify growth conditions with and without the ncAA supplied, respectively. Violet shades identify the cell populations collected. Arrows point to the following selection scheme applied to the collected cells after amplification by culturing.

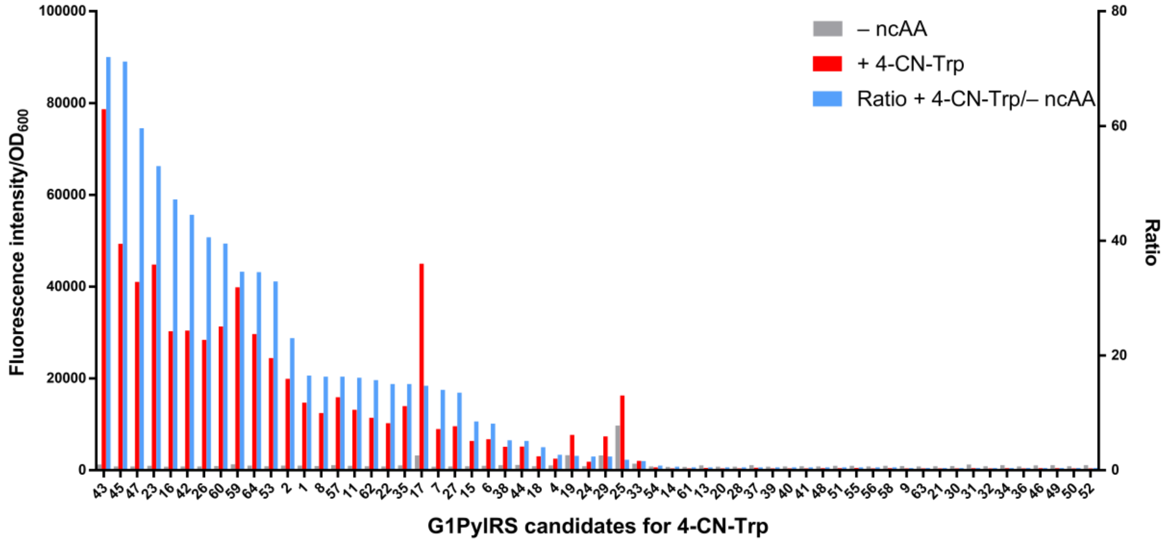

– ncAA
+ 4-CN-Trp
Ratio + 4-CN-Trp/– ncAA
Fluorescence intensity/OD600
Ratio
G1PylRS candidates for 4-CN-Trp

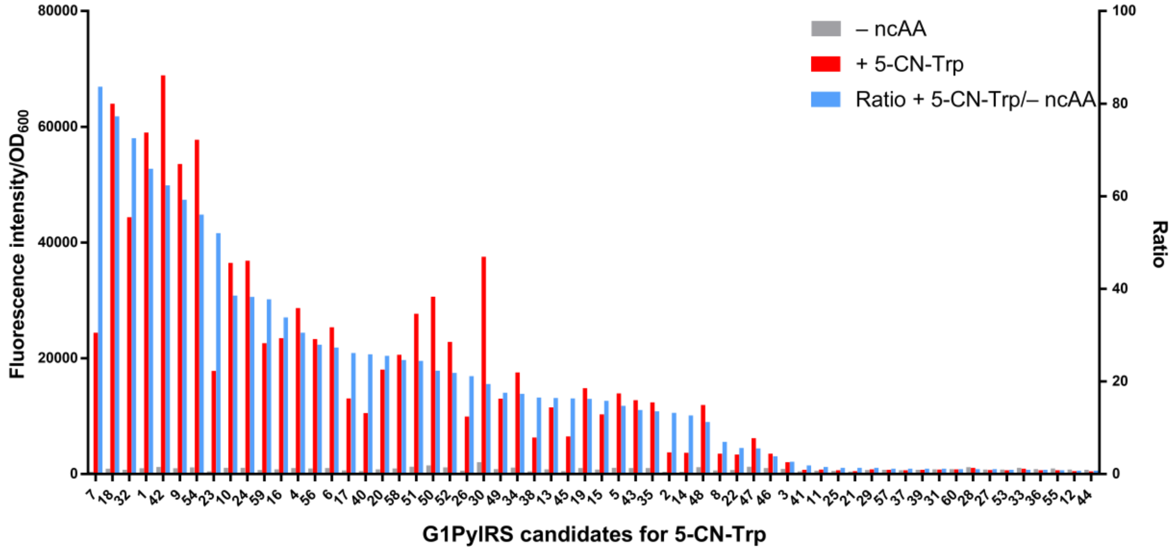

– ncAA
+ 5-CN-Trp
Ratio + 5-CN-Trp/– ncAA
Fluorescence intensity/OD600
Ratio
G1PylRS candidates for 5-CN-Trp

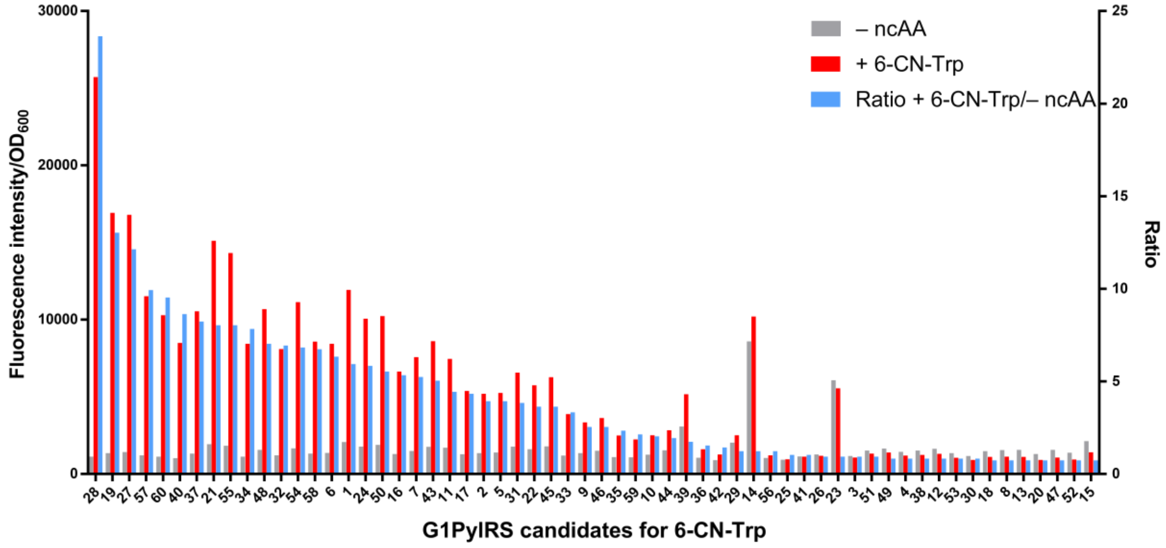

– ncAA
+ 6-CN-Trp
Ratio + 6-CN-Trp/– ncAA
Fluorescence intensity/OD600
Ratio
G1PylRS candidates for 6-CN-Trp

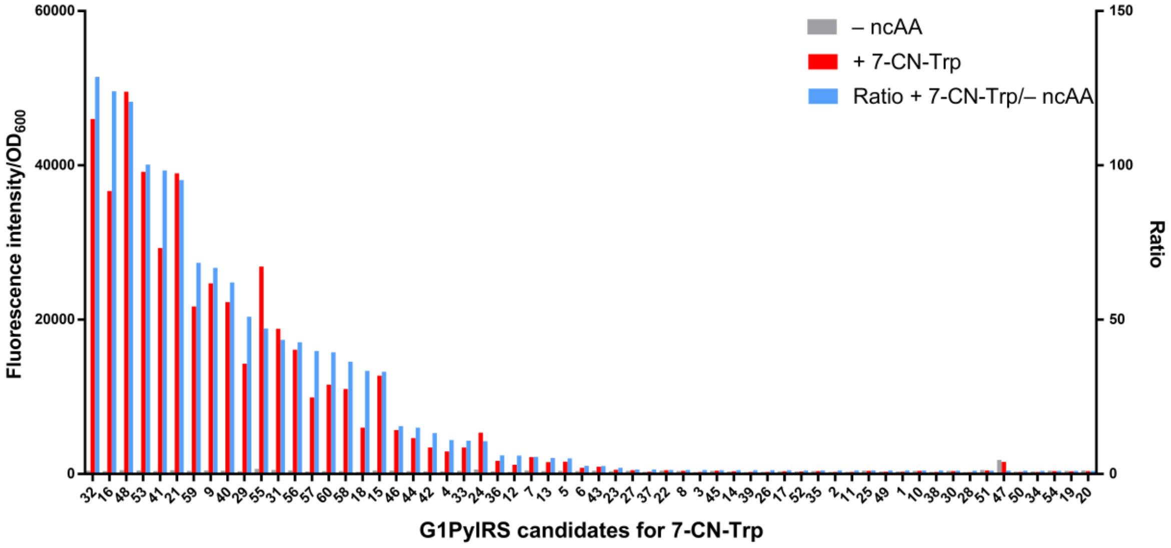

**Figure S5.** Activity and specificity screening of G1PylRS candidates for incorporating (a) 4-CN-Trp, (b) 5-CN-Trp, (c) 6-CN-Trp, or (d) 7-CN-Trp. The cells collected from the 3%–4% fraction with the highest red fluorescence in the final selection round were cultured in 96-well plates with and without 1 mM ncAA, and the intensity of red fluorescence was measured as an indicator of the readthrough efficiency of the amber-interrupted gene. The plot displays the colonies sorted by descending ratio of red fluorescence readouts from +ncAA wells relative to –ncAA wells.

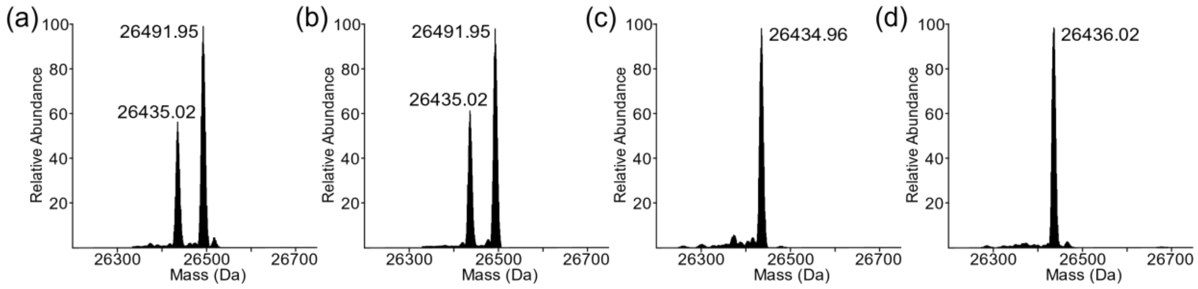

**Figure S6.** Intact protein mass spectrometry of His$_6$-TAG-mCherry with (a) 4-CN-Trp, (b) 5-CN-Trp, (c) 6-CN-Trp, or (d) 7-CN-Trp incorporated at the amber position. The calculated mass is 26434.79 Da. The +57 Da ions observed in the first two samples are a mass spectrometer artifact we often observe for mCherry RFP (but not GFP), possibly arising from iron leached out from the steel injector needle by the formic acid solvent used.

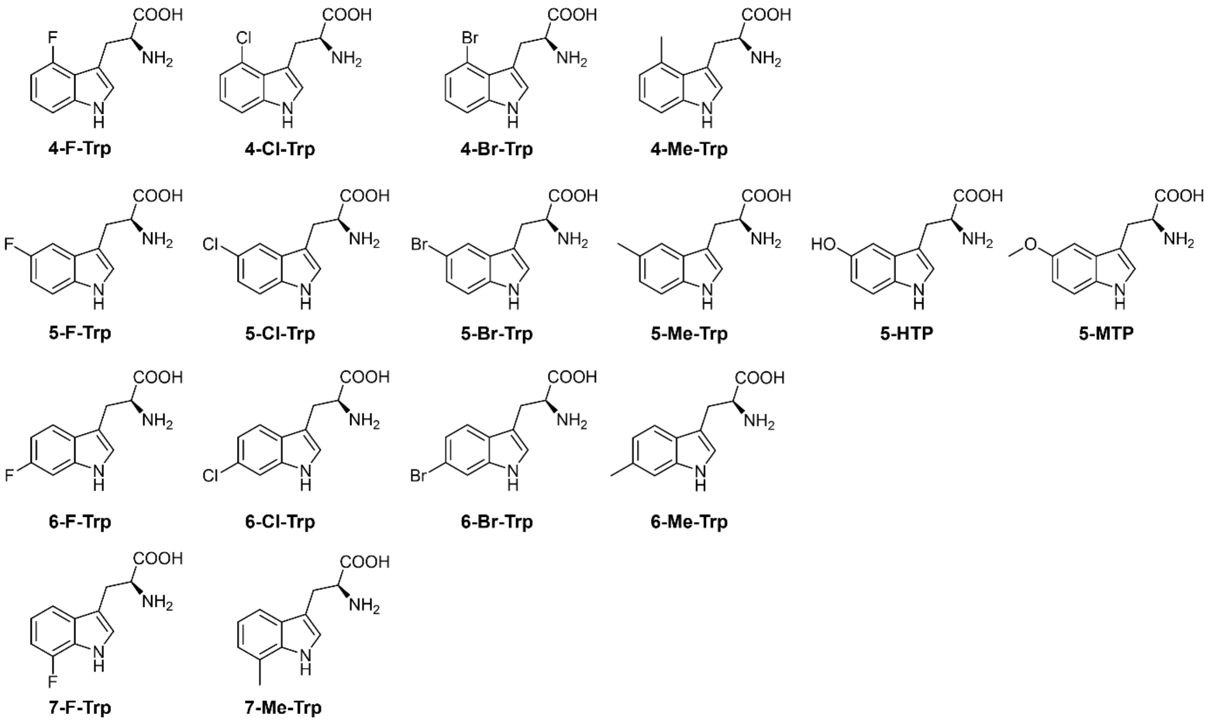

**Figure S7.** Chemical structures of other tryptophan-based non-canonical amino acids used in the present study.

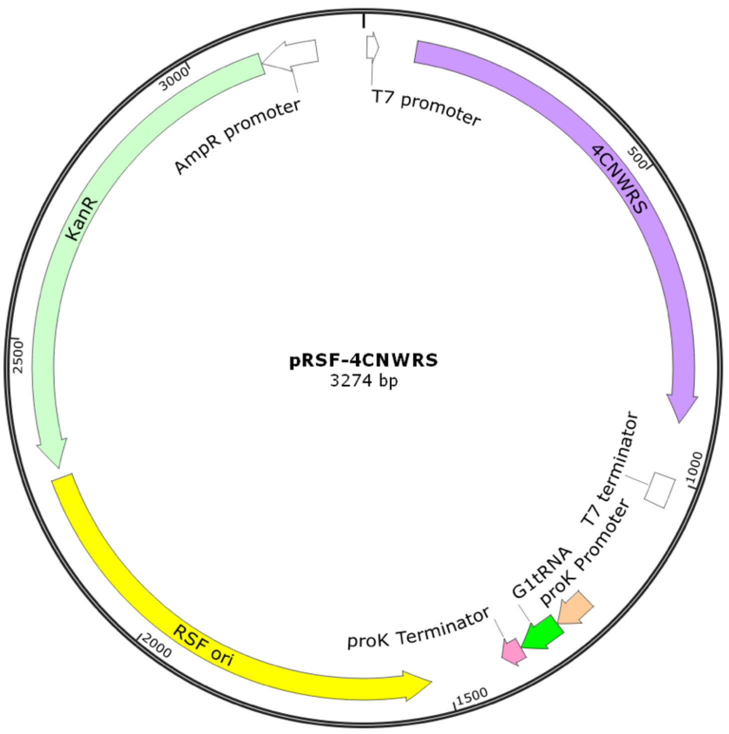

**Figure S8.** Plasmid map of pRSF-4CNWRS (Addgene No. #222871). The plasmid codes for the aminoacyl-tRNA synthetase and the requisite tRNA.

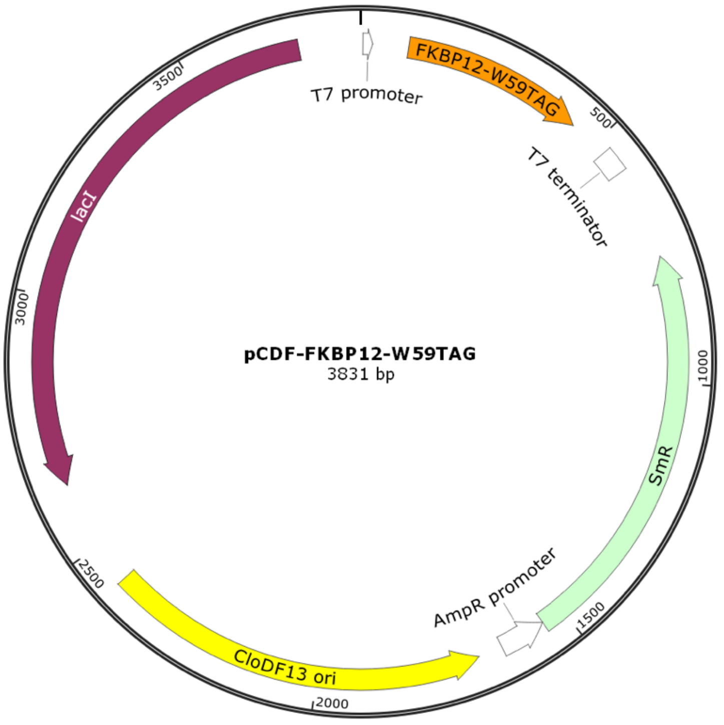

**Figure S9.** Plasmid map of pCDF-FKBP12-W59TAG. The plasmid codes for the target protein (FKBP12) with amber stop codon.

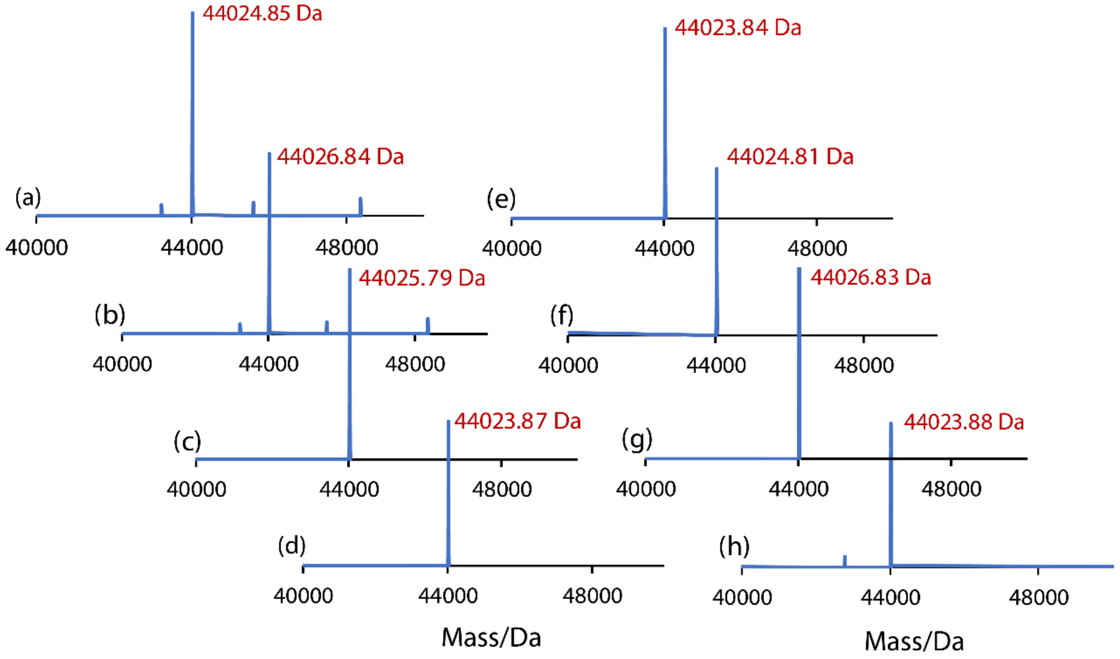

**Figure S10.** Intact protein mass spectrometry of mutants of the *E. coli* maltose binding protein, where individual tryptophan residues were replaced by 4-CN-Trp. The calculated mass of these mutants is 44,025.98 Da. The substituted tryptophan residues are (a) Trp10, (b) Trp62, (c) Trp94, (d) Trp129, (e) Trp158, (f) Trp230, (g) Trp232, (h) Trp340.

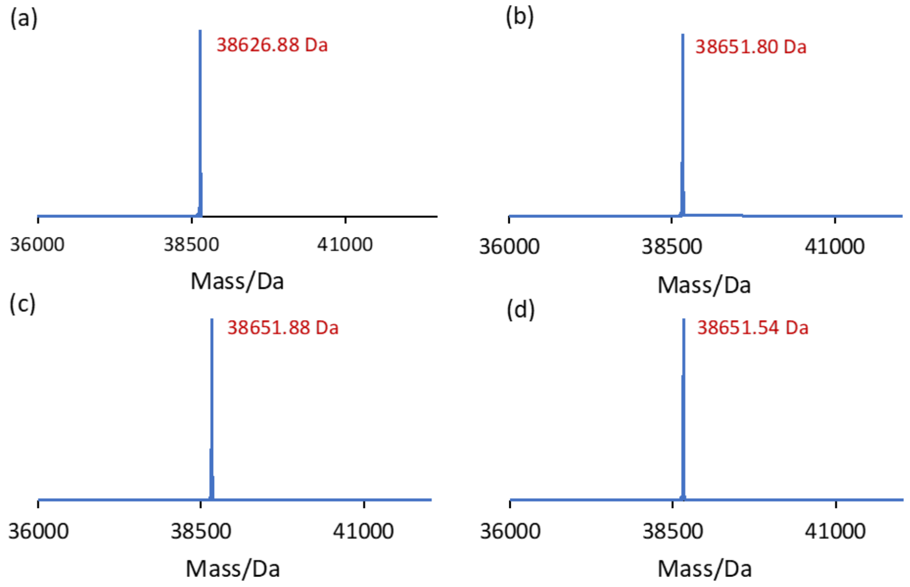

**Figure S11.** Mass spectra of hPXR-LBD with different tryptophan residues substituted by 4-CN-Trp. (a) Wild-type protein. The calculated mass is 38,628.49 Da. (b)–(d) Mutants with 4-CN-Trp installed in place of Trp199, Trp223 or Trp299, respectively. The calculated mass of these mutants is 38,653.50 Da.

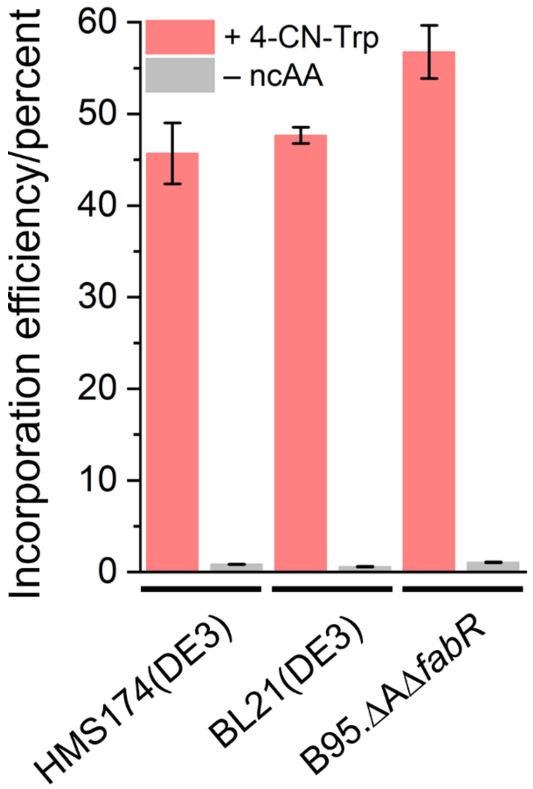

**Figure S12.** Efficiency of stop codon read-through by 4-CN-Trp in different *E. coli* strains. The cell lines HMS174(DE3), BL21(DE3) and B95.ΔAΔ*fabR* were transformed with both the pRSF-4CNWRS plasmid and the pCDF plasmid carrying the mCherry gene preceded by an amber stop codon. mCherry fluorescence was measured after overnight expression in 15 mL Falcon tubes with (red bars) or without (grey bars) the addition of 1 mM 4-CN-Trp and further normalized by the cell density. The incorporation efficiencies shown are relative to the fluorescence of the same mCherry construct expressed under the same conditions, except that the amber stop codon was replaced by a methionine codon.

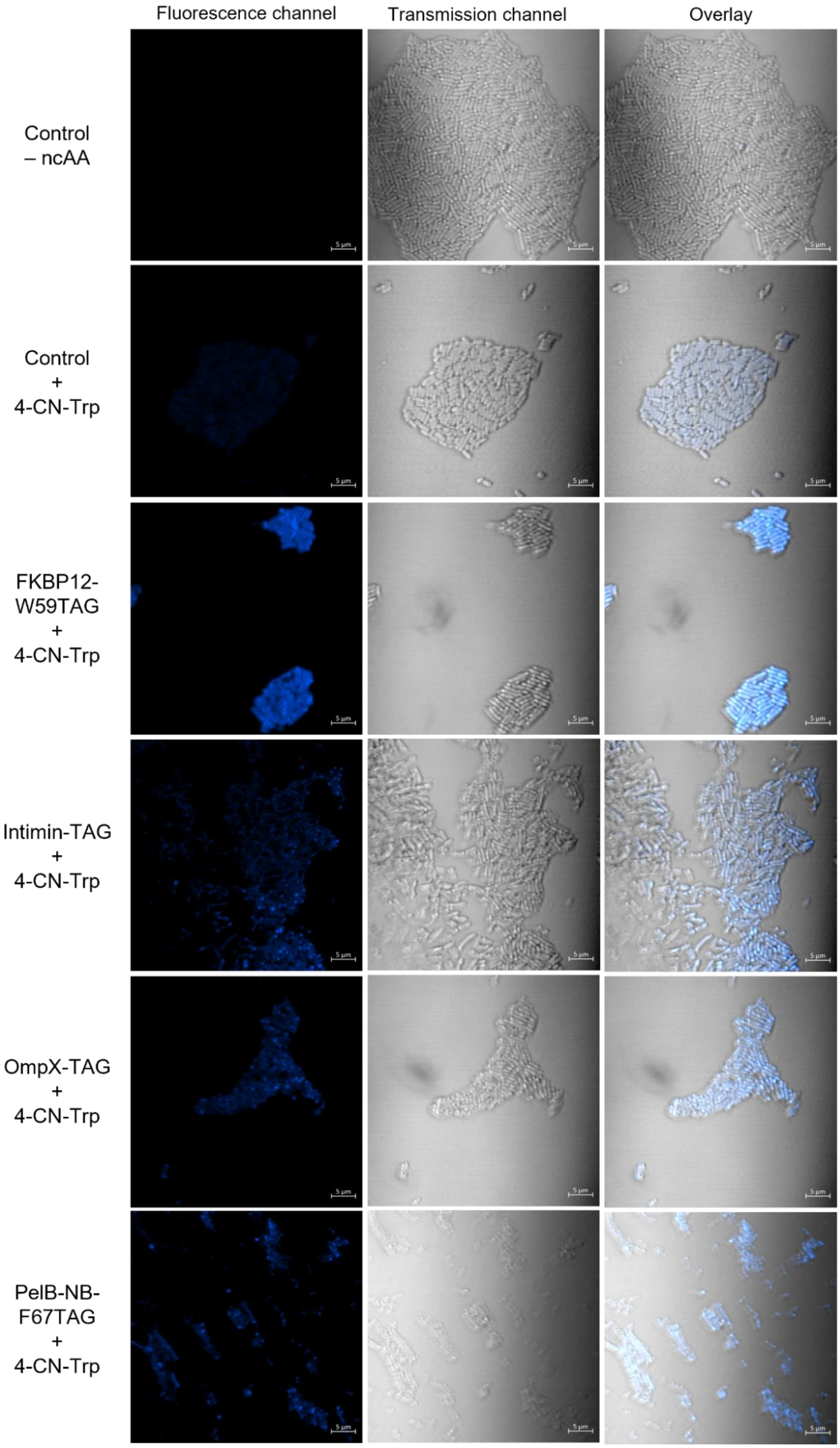
Fluorescence channel
Transmission channel
Overlay
Control
– ncAA
Control
+
4-CN-Trp
FKBP12-
W59TAG
+
4-CN-Trp
Intimin-TAG
+
4-CN-Trp
OmpX-TAG
+
4-CN-Trp
PelB-NB-
F67TAG
+
4-CN-Trp

**Figure S13.** Confocal fluorescence microscopy images of live *E. coli* cells expressing different 4-CN-Trp-labeled reporter proteins (scale bar = 5 μm). The samples labeled "+ 4-CN-Trp" were cultured in the presence of 0.75 mM 4-CN-Trp. The sample "Control + 4-CN-Trp" presents the control experiment for the baseline, where the cells lack the incorporation system for 4-CN-Trp. The images were taken using a 355 nm laser for excitation. The microscopy images were processed by combining the emission signals of all channels below 503 nm. The transmission channel was slightly out of focus when the focus was adjusted for the fluorescence channel, which may be attributed to UV light refraction by the agarose pad used as cell support.

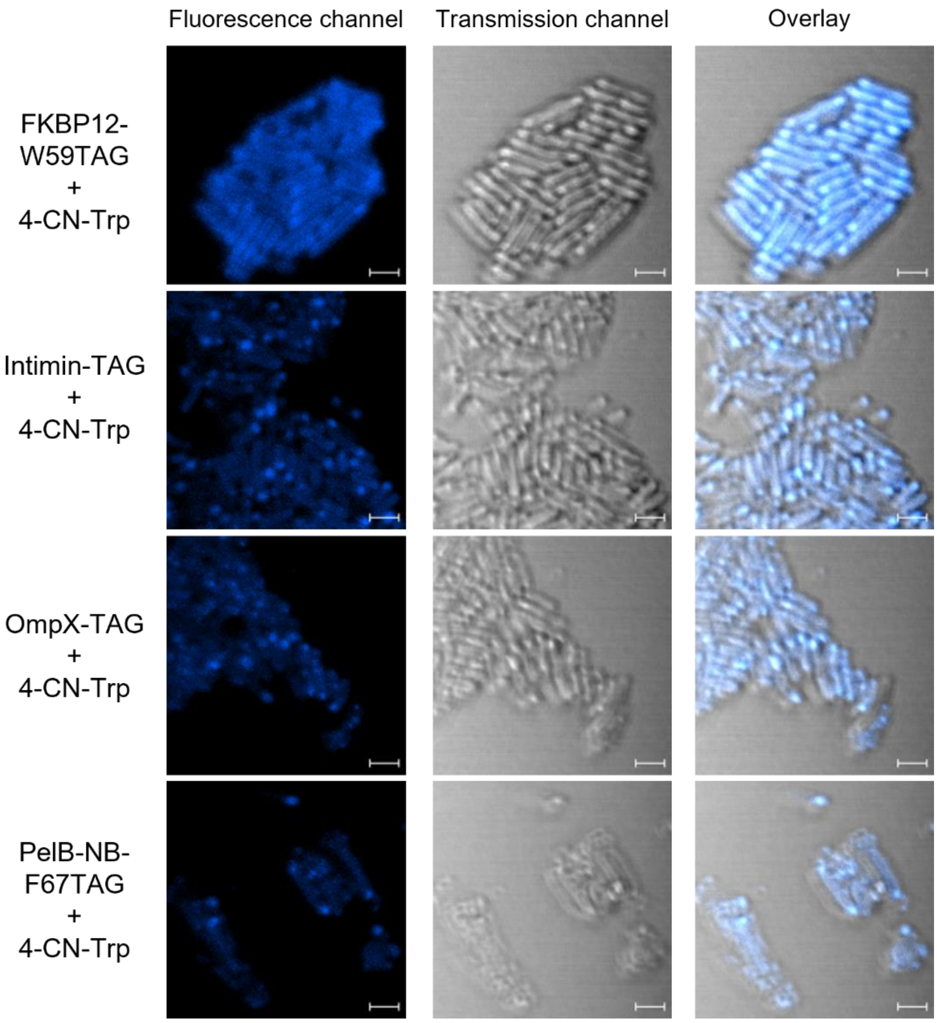

**Figure S14.** Selected regions from Figure S13 at increased magnification. Scale bar = 2 μm.

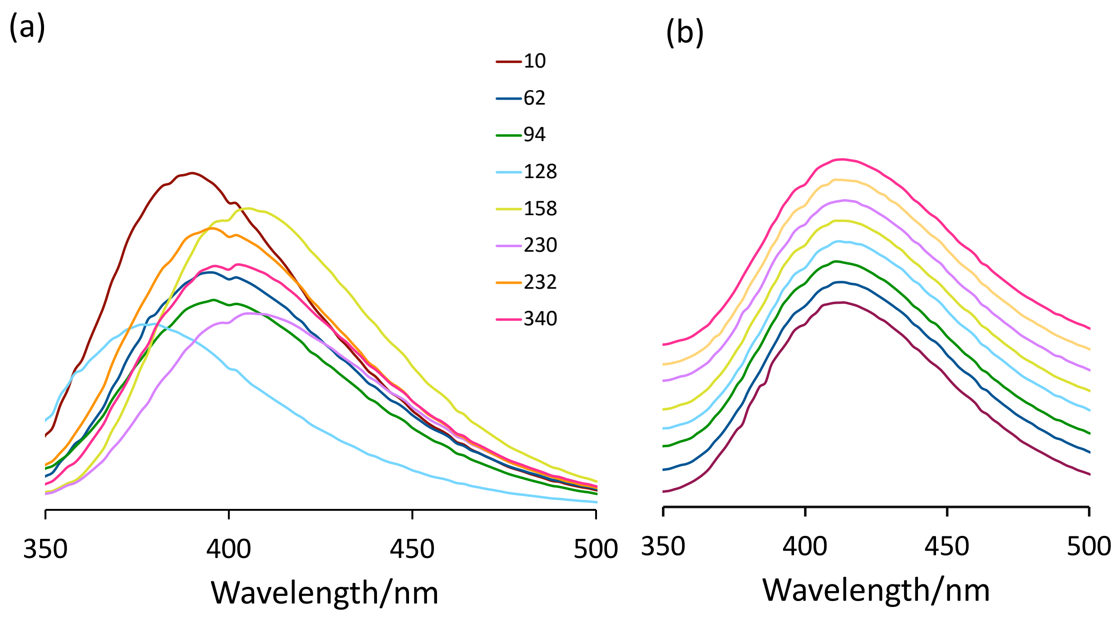

**Figure S15.** Fluorescence emission spectra of 1 μM solutions of MBP 4-CN-Trp mutants. The data were measured with an excitation wavelength of 310 nm on a TECAN Infinite 200 Pro M Plex plate reader (Tecan, Switzerland). (a) The emission spectra are color-coded with the sequence position of the Trp residue replaced by 4-CN-Trp as indicated. As the optical properties vary between different wells of the plate, the wavelengths of the emission maxima are accurate, but the intensities are unreliable. (b) Solutions in 8 M urea. To facilitate comparison, the emission spectra are scaled to the same height and vertically displayed relative to each other.

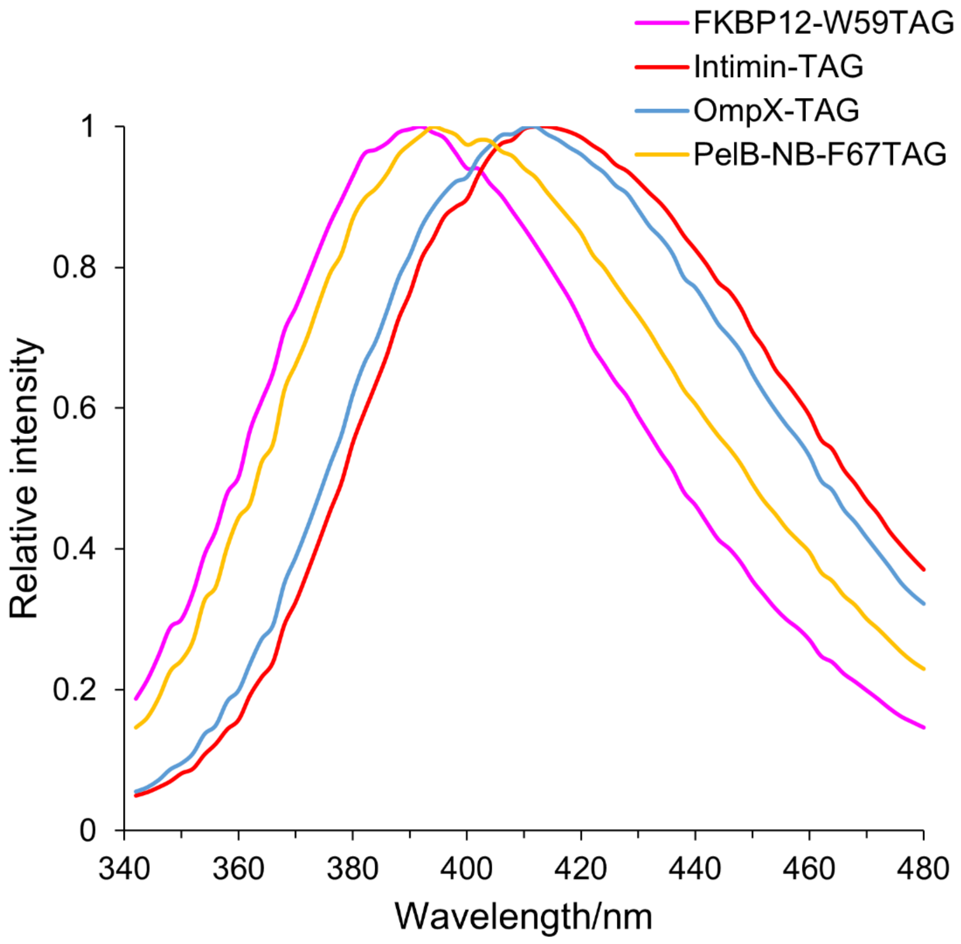

**Figure S16.** Fluorescence emission spectra of *E. coli* cells expressing various 4-CN-Trp-labeled reporter proteins measured with an excitation wavelength of 310 nm on a TECAN Infinite 200 Pro M Plex plate reader (Tecan, Switzerland). The fluorescence intensities are normalized by the maximum value of each curve.

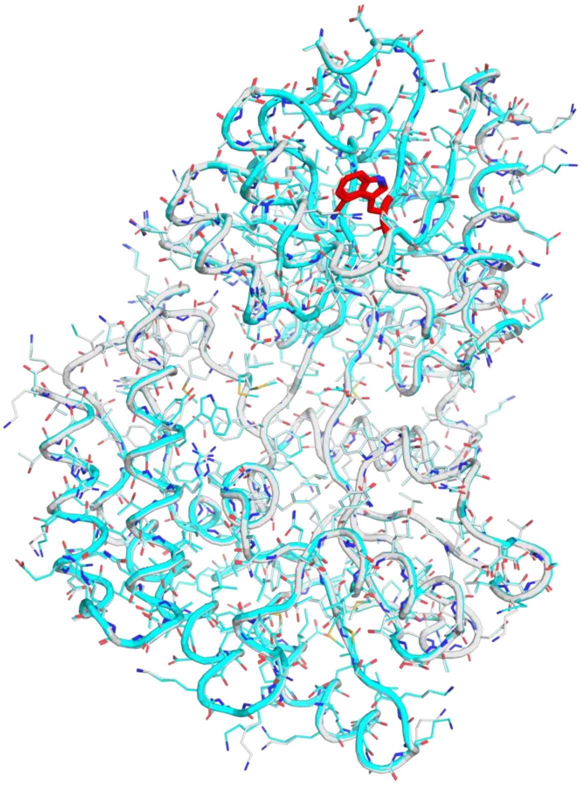

**Figure S17.** Superimposition of the crystal structures of maltose binding protein (MBP), wild-type (grey) and produced with 4-CN-Trp in position 10 (cyan), illustrating the level of structural conservation. The backbone atoms superimpose with a root mean square deviation of < 0.1 Å. The residue in position 10 is highlighted in red.

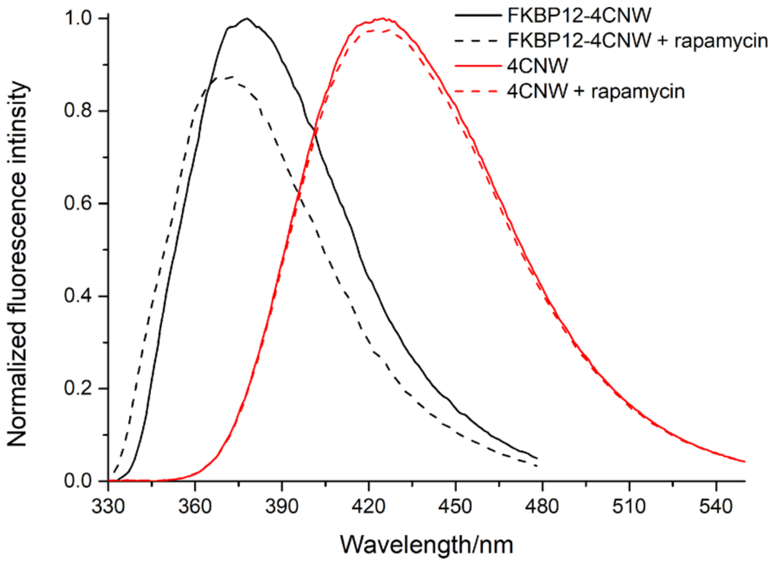

**Figure S18.** Attenuation of 4-CN-Trp fluorescence by rapamycin. The black curves show the fluorescence emission of a 1 μM solution of FKBP12-4CNW in the absence (solid line) and presence (dashed line) of a 20-fold excess of rapamycin. These data are the same as those shown in Figure 5. The red curves show the corresponding data obtained with a 1 μM solution of 4-CN-Trp instead of FKBP12-4CNW.

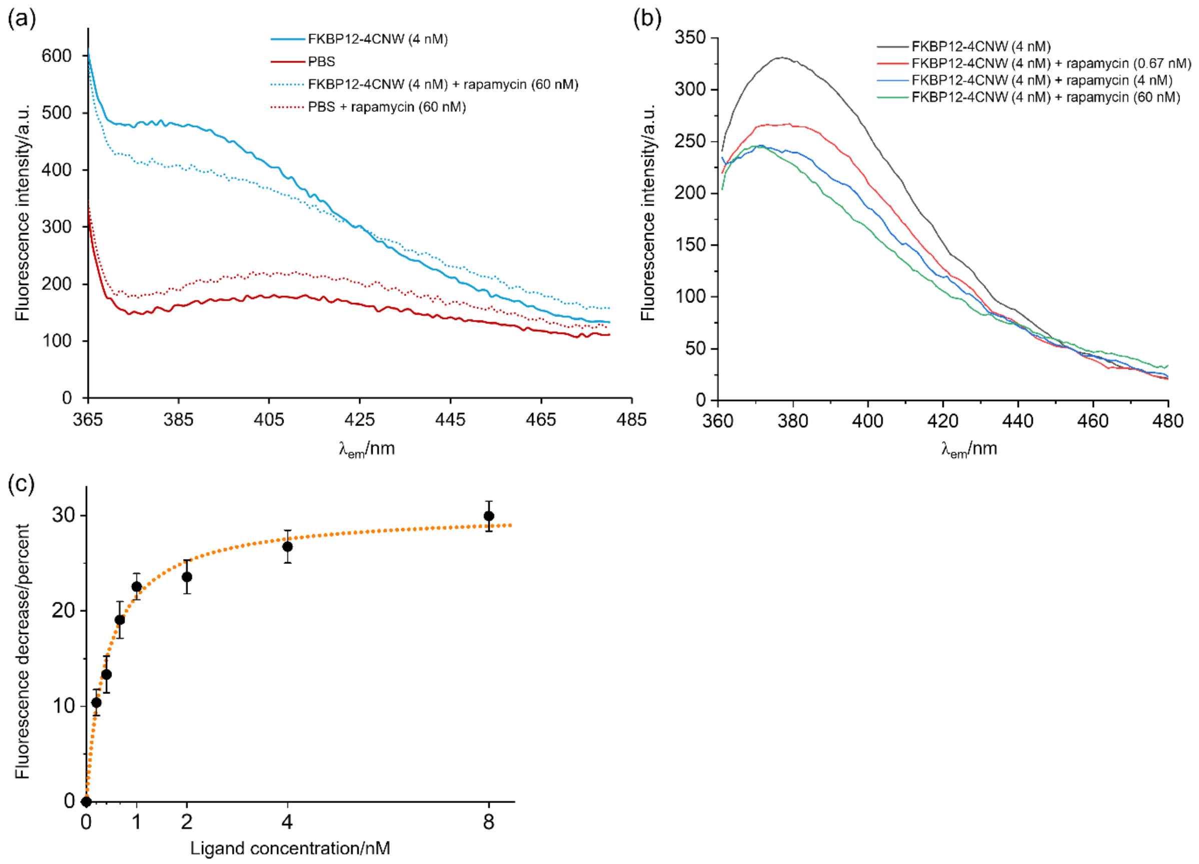

**Figure S19.** Fluorescence emission spectra of a 4 nM solution of FKBP12-4CNW recorded at room temperature in PBS buffer with and without the presence of rapamycin. The measurements were carried out in a 1 cm quartz cuvette using 95% power of the excitation source (excitation wavelength 310 nm). For improved sensitivity, the emission intensity at 380 nm was calculated by averaging five data points between 378 nm and 382 nm. (a) Raw data and background measurements. Background data were measured with buffer only and buffer plus rapamycin only for subtraction from the signals measured with FKBP12-4CNW and FKBP12-4CNW plus rapamycin. Note that the fluorescence of rapamycin contributes to the background fluorescence, whereas the presence of rapamycin in the protein sample decreases the overall fluorescence intensity. (b) Spectra after background subtraction and smoothing by averaging over five data points separated by 1 nm. (c) Zoomed-in view of the titration curve at low concentration range of Figure b.

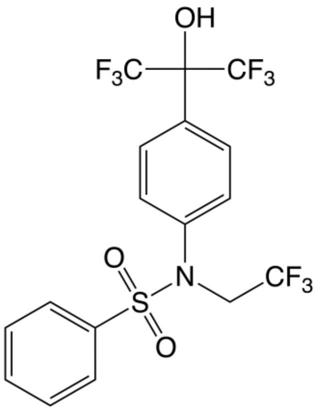

**Figure S20.** Chemical structure of TO901317 (compound **1**).

#### Supplementary tables

**Table S1.** Fluorescence characteristics of CN-tryptophans measured with the samples of Figure S2.

| Amino acid | $\lambda_{ex}$ (nm) | $\lambda_{em}$ (nm) | Extinction coefficient[a] ($M^{-1}$ $cm^{-1}$) | Quantum yield (%)[b] | Brightness[c] ($M^{-1}$ $cm^{-1}$) |
|---|---|---|---|---|---|
| 4-CN-Trp | 305 | 424 | 6041 ± 58 (at 305 nm) | 72 ± 2 | 4300 |
| | | | 2304 ± 48 (at 280 nm) | | 1700 |
| 5-CN-Trp | 280 | - | 4835 ± 78 (at 280 nm) | < 1 | - |
| 6-CN-Trp | 280 | 380 | 5940 ± 74 (at 280 nm) | 23.1 ± 0.2 | 1400 |
| 7-CN-Trp | 305 | 416 | 8289 ± 70 (at 305 nm) | 12.6 ± 0.2 | 1000 |
| | | | 3064 ± 43 (at 280 nm) | | 400 |

[a] Extinction coefficients at 280 nm are useful for determining protein concentrations based on their amino acid sequence. For comparison, the $\varepsilon_{280}$ value of tryptophan is 5422 $M^{-1}$ $cm^{-1}$.

[b] The quantum yield was determined using the protocol of Würth et al.[22] using either tryptophan (280 nm) or quinine sulfate (305 nm) as reference standard.

[c] Calculated as the product of extinction coefficient and quantum yield.

**Table S2.** Mutations found in twelve G1PylRS variants selected for activity with 4-CN-Trp, 5-CN-Trp, 6-CN-Trp, or 7-CN-Trp.

| **RS Variants** | **Randomized Sites** | | | | | | |
|---|---|---|---|---|---|---|---|
| *Mm*PylRS wild-type[a] | L305 | Y306 | N346 | V348 | Y384 | V401 | W417 |
| *G1*PylRS wild-type | L124 | Y125 | N165 | V167 | Y204 | A221 | W237 |
| 4CNW16 | L | Y | A | S | W | S | G |
| 4CNW45[b] | I | F | G | A | Y | T | S |
| 4CNW47 | L | A | A | C | W | S | H |
| 4CNW59 | L | Y | A | F | W | S | A |
| 5CNW10 | V | F | G | V | W | S | V |
| 5CNW32[b] | V | F | G | V | F | S | S |
| 5CNW42 | T | F | G | C | W | T | T |
| 6CNW27 | L | Y | A | A | Y | G | S |
| 6CNW28[b] | H | Y | T | G | Y | G | S |
| 7CNW09 | Q | F | G | S | W | G | Y |
| 7CNW16 | L | M | A | F | W | G | K |
| 7CNW32[b] | A | F | G | G | W | G | Y |

[a] Corresponding sites in the *Methanosarcina mazei* PylRS enzyme provided for comparison.

[b] The mutants 4CNW45, 5CNW32, 6CNW28, and 7CNW32 were used for all following applications and named 4CNWRS, 5CNWRS, 6CNWRS, and 7CNWRS, respectively.

**Table S3.** DNA and corresponding amino acid sequences of the proteins used in the current study.

| Protein | DNA sequence | Amino acid sequence[a] |
|---|---|---|
| 4CNWRS | ATGGTGGTGAAATTTACCGATAGCCAGATTCAGCATCTGATGGAATA<br>TGGTGATAATGATTGGAGCGAAGCCGAATTTGAAGATGCAGCAGCAC<br>GTGATAAAGAATTTAGCAGCCAGTTTAGCAAACTGAAAAGCGCCAAT<br>GATAAAGGCCTGAAAGATGTTATTGCAAATCCGCGTAATGATCTGAC<br>CGATCTGGAAAACAAAATTCGCGAAAAACTGGCAGCCCGTGGTTTTA<br>TTGAAGTTCATACCCCGATTTTTGTGAGCAAAAGCGCACTGGCAAAA<br>ATGACCATTACCGAAGATCATCCGCTGTTCAAACAGGTGTTTTGGAT<br>TGATGATAAACGTGCACTGCGTCCGATGCATGCAATGAATATTTTTA<br>AAGTTATGCGTGAACTGCGCGATCATACCAAAGGTCCGGTTAAAATC<br>TTTGAAATTGGTAGCTGCTTTCGCAAAGAAAGCAAAAGCAGTACCCA<br>TCTGGAAGAATTTACCATGCTGGGCCTGGCCGAAATGGGTCCTGATG<br>GTGATCCGATGGAACATCTGAAAATGTATATTGGCGATATCATGGAT<br>GCCGTTGGTGTTGAATATACCACCAGTCGTGAAGAATCAGATGTTTA<br>TGTTGAAACCCTGGACGTGGAAATTAATGGCACCGAAGTTGCAAGCG<br>GTACTGTTGGTCCGCATAAACTGGATCCGGCACATGATGTGCATGAA<br>CCGTCGGCAGGTATTGGTTTTGGTCTGGAACGTCTGCTGATGCTGAA<br>AAATGGTAAAAGCAATGCACGCAAAACCGGCAAAAGTATTACCTATC<br>TGAATGGCTACAAACTGGATTAA | MVVKFTDSQIQHLMEYGDNDWSE<br>AEFEDAAARDKEFSSQFSKLKSA<br>NDKGLKDVIANPRNDLTDLENKI<br>REKLAARGFIEVHTPIFVSKSAL<br>AKMTITEDHPLFKQVFWIDDKRA<br>LRPMHAMNIFKVMRELRDHTKGP<br>VKIFEIGSCFRKESKSSTHLEEF<br>TMLGLAEMGPDGDPMEHLKMYIG<br>DIMDAVGVEYTTSREESDVYVET<br>LDVEINGTEVASGTVGPHKLDPA<br>HDVHEPSAGIGFGLERLLMLKNG<br>KSNARKTGKSITYLNGYKLD |
| 5CNWRS | ATGGTGGTGAAATTTACCGATAGCCAGATTCAGCATCTGATGGAATA<br>TGGTGATAATGATTGGAGCGAAGCCGAATTTGAAGATGCAGCAGCAC<br>GTGATAAAGAATTTAGCAGCCAGTTTAGCAAACTGAAAAGCGCCAAT<br>GATAAAGGCCTGAAAGATGTTATTGCAAATCCGCGTAATGATCTGAC<br>CGATCTGGAAAACAAAATTCGCGAAAAACTGGCAGCCCGTGGTTTTA<br>TTGAAGTTCATACCCCGATTTTTGTGAGCAAAAGCGCACTGGCAAAA<br>ATGACCATTACCGAAGATCATCCGCTGTTCAAACAGGTGTTTTGGAT<br>TGATGATAAACGTGCACTGCGTCCGATGCATGCAATGAATGTGTTTA<br>AAGTTATGCGTGAACTGCGCGATCATACCAAAGGTCCGGTTAAAATC<br>TTTGAAATTGGTAGCTGCTTTCGCAAAGAAAGCAAAAGCAGTACCCA<br>TCTGGAAGAATTTACCATGCTGGGCCTGGTAGAAATGGGTCCTGATG<br>GTGATCCGATGGAACATCTGAAAATGTATATTGGCGATATCATGGAT<br>GCCGTTGGTGTTGAATATACCACCAGTCGTGAAGAATCAGATGTTTT<br>TGTTGAAACCCTGGACGTGGAAATTAATGGCACCGAAGTTGCAAGCG<br>GTTCTGTTGGTCCGCATAAACTGGATCCGGCACATGATGTGCATGAA<br>CCGAGTGCAGGTATTGGTTTTGGTCTGGAACGTCTGCTGATGCTGAA<br>AAATGGTAAAAGCAATGCACGCAAAACCGGCAAAAGTATTACCTATC<br>TGAATGGCTACAAACTGGATTAA | MVVKFTDSQIQHLMEYGDNDWSE<br>AEFEDAAARDKEFSSQFSKLKSA<br>NDKGLKDVIANPRNDLTDLENKI<br>REKLAARGFIEVHTPIFVSKSAL<br>AKMTITEDHPLFKQVFWIDDKRA<br>LRPMHAMNVFKVMRELRDHTKGP<br>VKIFEIGSCFRKESKSSTHLEEF<br>TMLGLVEMGPDGDPMEHLKMYIG<br>DIMDAVGVEYTTSREESDVFVET<br>LDVEINGTEVASGSVGPHKLDPA<br>HDVHEPSAGIGFGLERLLMLKNG<br>KSNARKTGKSITYLNGYKLD |
| 6CNWRS | ATGGTGGTGAAATTTACCGATAGCCAGATTCAGCATCTGATGGAATA<br>TGGTGATAATGATTGGAGCGAAGCCGAATTTGAAGATGCAGCAGCAC<br>GTGATAAAGAATTTAGCAGCCAGTTTAGCAAACTGAAAAGCGCCAAT<br>GATAAAGGCCTGAAAGATGTTATTGCAAATCCGCGTAATGATCTGAC<br>CGATCTGGAAAACAAAATTCGCGAAAAACTGGCAGCCCGTGGTTTTA<br>TTGAAGTTCATACCCCGATTTTTGTGAGCAAAAGCGCACTGGCAAAA<br>ATGACCATTACCGAAGATCATCCGCTGTTCAAACAGGTGTTTTGGAT<br>TGATGATAAACGTGCACTGCGTCCGATGCATGCAATGAATCATTATA<br>AAGTTATGCGTGAACTGCGCGATCATACCAAAGGTCCGGTTAAAATC<br>TTTGAAATTGGTAGCTGCTTTCGCAAAGAAAGCAAAAGCAGTACCCA<br>TCTGGAAGAATTTACCATGCTGACCCTGGGAGAAATGGGTCCTGATG<br>GTGATCCGATGGAACATCTGAAAATGTATATTGGCGATATCATGGAT<br>GCCGTTGGTGTTGAATATACCACCAGTCGTGAAGAATCAGATGTTTA<br>TGTTGAAACCCTGGACGTGGAAATTAATGGCACCGAAGTTGCAAGCG<br>GTGGTGTTGGTCCGCATAAACTGGATCCGGCACATGATGTGCATGAA<br>CCGAGTGCAGGTATTGGTTTTGGTCTGGAACGTCTGCTGATGCTGAA<br>AAATGGTAAAAGCAATGCACGCAAAACCGGCAAAAGTATTACCTATC<br>TGAATGGCTACAAACTGGATTAA | MVVKFTDSQIQHLMEYGDNDWSE<br>AEFEDAAARDKEFSSQFSKLKSA<br>NDKGLKDVIANPRNDLTDLENKI<br>REKLAARGFIEVHTPIFVSKSAL<br>AKMTITEDHPLFKQVFWIDDKRA<br>LRPMHAMNHYKVMRELRDHTKGP<br>VKIFEIGSCFRKESKSSTHLEEF<br>TMLTLGEMGPDGDPMEHLKMYIG<br>DIMDAVGVEYTTSREESDVYVET<br>LDVEINGTEVASGGVGPHKLDPA<br>HDVHEPSAGIGFGLERLLMLKNG<br>KSNARKTGKSITYLNGYKLD |

| | | |
|---|---|---|
| 7CNWRS | ATGGTGGTGAAATTTACCGATAGCCAGATTCAGCATCTGATGGAATA<br>TGGTGATAATGATTGGAGCGAAGCCGAATTTGAAGATGCAGCAGCAC<br>GTGATAAAGAATTTAGCAGCCAGTTTAGCAAACTGAAAAGCGCCAAT<br>GATAAAGGCCTGAAAGATGTTATTGCAAATCCGCGTAATGATCTGAC<br>CGATCTGGAAAACAAAATTCGCGAAAAACTGGCAGCCCGTGGTTTTA<br>TTGAAGTTCATACCCCGATTTTTGTGAGCAAAAGCGCACTGGCAAAA<br>ATGACCATTACCGAAGATCATCCGCTGTTCAAACAGGTGTTTTGGAT<br>TGATGATAAACGTGCACTGCGTCCGATGCATGCAATGAATGCTTTTA<br>AAGTTATGCGTGAACTGCGCGATCATACCAAAGGTCCGGTTAAAATC<br>TTTGAAATTGGTAGCTGCTTTCGCAAAGAAAGCAAAAGCAGTACCCA<br>TCTGGAAGAATTTACCATGCTGGGCCTGGGAGAAATGGGTCCTGATG<br>GTGATCCGATGGAACATCTGAAAATGTATATTGGCGATATCATGGAT<br>GCCGTTGGTGTTGAATATACCACCAGTCGTGAAGAATCAGATGTTTG<br>GGTTGAAACCCTGGACGTGGAAATTAATGGCACCGAAGTTGCAAGCG<br>GTGGGGTTGGTCCGCATAAACTGGATCCGGCACATGATGTGCATGAA<br>CCGTATGCAGGTATTGGTTTTGGTCTGGAACGTCTGCTGATGCTGAA<br>AAATGGTAAAAGCAATGCACGCAAAACCGGCAAAAGTATTACCTATC<br>TGAATGGCTACAAACTGGATTAA | MVVKFTDSQIQHLMEYGDNDWSE<br>AEFEDAAARDKEFSSQFSKLKSA<br>NDKGLKDVIANPRNDLTDLENKI<br>REKLAARGFIEVHTPIFVSKSAL<br>AKMTITEDHPLFKQVFWIDDKRA<br>LRPMHAMNAFKVMRELRDHTKGP<br>VKIFEIGSCFRKESKSSTHLEEF<br>TMLGLGEMGPDGDPMEHLKMYIG<br>DIMDAVGVEYTTSREESDVWVET<br>LDVEINGTEVASGGVGPHKLDPA<br>HDVHEPYAGIGFGLERLLMLKNG<br>KSNARKTGKSITYLNGYKLD |
| $His_6$-TAG-RFP | ATGCACCACCATCACCATCACTAGGCCAGTAGTGAAGACGTTATCAA<br>GGAGTTTATGCGTTTCAAAGTACGTATGGAGGGTAGTGTTAACGGAC<br>ACGAATTTGAGATCGAGGGAGAGGGGGAAGGTCGTCCTTACGAGGGA<br>ACTCAAACGGCCAAATTAAAGGTGACCAAAGGTGGGCCCTTGCCATT<br>CGCGTGGGACATCTTGTCACCCCAGTTCCAGTACGGGTCGAAGGCAT<br>ACGTAAAACACCCAGCGGACATTCCTGACTATCTTAAGTTATCTTTC<br>CCGGAAGGTTTTAAATGGGAACGCGTGATGAACTTTGAGGATGGGGG<br>GGTTGTTACGGTGACACAAGACTCCTCATTGCAAGATGGAGAGTTTA<br>TCTATAAAGTCAAACTTCGCGGCACCAATTTTCCATCTGACGGTCCT<br>GTAATGCAGAAAAAAACAATGGGCTGGGAAGCCTCCACAGAACGTAT<br>GTACCCCGAAGATGGAGCTTTAAAGGGCGAAATTAAAATGCGCTTAA<br>AACTTAAAGACGGCGGCCATTACGACGCCGAAGTGAAAACGACGTAT<br>ATGGCTAAGAAACCCGTCCAGCTTCCGGGAGCCTATAAAACTGACAT<br>CAAACTGGATATTACATCACACAACGAAGATTATACTATTGTCGAAC<br>AGTACGAACGCGCCGAAGGCCGCCATTCAACGGGAGCATAA | MHHHHHHXASSEDVIKEFMRFKV<br>RMEGSVNGHEFEIEGEGEGRPYE<br>GTQTAKLKVTKGGPLPFAWDILS<br>PQFQYGSKAYVKHPADIPDYLKL<br>SFPEGFKWERVMNFEDGGVVTVT<br>QDSSLQDGEFIYKVKLRGTNFPS<br>DGPVMQKKTMGWEASTERMYPED<br>GALKGEIKMRLKLKDGGHYDAEV<br>KTTYMAKKPVQLPGAYKTDIKLD<br>ITSHNEDYTIVEQYERAEGRHST<br>GA |
| FKBP12-<br>W59TAG | ATGGGTGTTCAGGTTGAAACCATTAGTCCTGGTGATGGTCGTACCTT<br>TCCGAAACGTGGTCAGACCTGTGTTGTTCATTACACCGGTATGCTGG<br>AAGATGGCAAAAAGTTTGATAGCAGCCGTGATCGTAATAAGCCGTTT<br>AAATTCGTTCTGGGTAAACAAGAAGTTATTCGCGGTTAGGAAGAGGG<br>TGTTGCACAGATGAGCGTTGGTCAGCGTGCAAAACTGACCATTTCAC<br>CGGATTATGCATATGGTGCAACCGGTCATCCGGGTATTATTCCGCCT<br>AATGCAACCCTGATTTTTGATGTTGAACTGCTGAAACTGGAAGGTGG<br>TAGCCATCACCATCATCATCATTAA | MGVQVETISPGDGRTFPKRGQTC<br>VVHYTGMLEDGKKFDSSRDRNKP<br>FKFVLGKQEVIRGXEEGVAQMSV<br>GQRAKLTISPDYAYGATGHPGII<br>PPNATLIFDVELLKLEGGSHHHH<br>HH |
| Intimin-TAG | ATGATTACTCATGGTTGTTATACCCGGACCCGGCACAAGCATAAGCT<br>AAAAAAAACATTGATTATGCTTAGTGCTGGTTTAGGATTGTTTTTTT<br>ATGTTAATCAGAACTCATTTGCAAATGGTGAAAATTATTTTAAATTG<br>GGTTCGGATTCAAAACTGTTAACTCATGATAGCTATCAGAATCGCCT<br>TTTTTATACGTTGAAAACTGGTGAAACTGTTGCCGATCTTTCTAAAT<br>CGCAAGATATTAATTTATCGACGATTTGGTCGTTGAATAAGCATTTA<br>TACAGTTCTGAAAGCGAAATGATGAAGGCCGCGCCTGGTCAGCAGAT<br>CATTTTGCCACTCAAAAAACTTCCCTTTGAATACAGTGCACTACCAC<br>TTTTAGGTTCGGCACCTCTTGTTGCTGCGGGTGGTGTTGCTGGTCAC<br>ACGAATAAACTGACTAAAATGTCCCCGGACGTGACCAAAAGCAACAT<br>GACCGATGACAAGGCATTAAATTATGCGGCACAACAGGCGGCGAGTC<br>TCGGTAGCCAGCTTCAGTCGCGATCTCTGAACGGCGATTACGCGAAA<br>GATACCGCTCTTGGTATCGCTGGTAACCAGGCTTCGTCACAGTTGCA<br>GGCCTGGTTACAACATTATGGAACGGCAGAGGTTAATCTGCAAAGTG<br>GTAATAACTTTGACGGTAGTTCACTGGACTTCTTATTACCGTTCTAT<br>GATTCCGAAAAAATGCTGGCATTTGGTCAGGTCGGAGCGCGTTACAT<br>TGACTCCCGCTTTACGGCAAATTTAGGTGCGGGTCAGCGTTTTTTCC<br>TTCCTGCAAACATGTTGGGCTATAACGTCTTCATTGATCAGGATTTT<br>TCTGGTGATAATACCCGTTTAGGTATTGGTGGCGAATACTGGCGAGA<br>CTATTTCAAAAGTAGCGTTAACGGCTATTTCCGCATGAGCGGCTGGC<br>ATGAGTCATACAATAAGAAAGACTATGATGAGCGCCCAGCAAATGGC | MITHGCYTRTRHKHKLKKTLIML<br>SAGLGLFFYVNQNSFANGENYFK<br>LGSDSKLLTHDSYQNRLFYTLKT<br>GETVADLSKSQDINLSTIWSLNK<br>HLYSSESEMMKAAPGQQIILPLK<br>KLPFEYSALPLLGSAPLVAAGGV<br>AGHTNKLTKMSPDVTKSNMTDDK<br>ALNYAAQQAASLGSQLQSRSLNG<br>DYAKDTALGIAGNQASSQLQAWL<br>QHYGTAEVNLQSGNNFDGSSLDF<br>LLPFYDSEKMLAFGQVGARYIDS<br>RFTANLGAGQRFFLPANMLGYNV<br>FIDQDFSGDNTRLGIGGEYWRDY<br>FKSSVNGYFRMSGWHESYNKKDY<br>DERPANGFDIRFNGYLPSYPALG<br>AKLIYEQYYGDNVALFNSDKLQS<br>NPGAATVGVNYTPIPLVTMGIDY<br>RHGTGNENDLLYSMQFRYQFDKS<br>WSQQIEPQYVNELRTLSGSRYDL<br>VQRNNNIILEYKKQDILSLNIPH<br>DINGTEHSTQKIQLIVKSKYGLD |

| | | |
|---|---|---|
| | TTCGATATCCGTTTTAATGGCTATCTACCGTCATATCCGGCATTAGG<br>CGCCAAGCTGATATATGAGCAGTATTATGGTGATAATGTTGCTTTGT<br>TTAATTCTGATAAGCTGCAATCGAATCCTGGTGCGGCGACCGTTGGT<br>GTAAACTATACTCCGATTCCTCTGGTGACGATGGGGATCGATTACCG<br>TCATGGTACGGGTAATGAAAATGATCTCCTTTACTCAATGCAGTTCC<br>GTTATCAGTTTGATAAATCGTGGTCTCAGCAAATTGAACCACAGTAT<br>GTTAACGAGTTAAGAACATTATCAGGCAGCCGTTACGATCTGGTTCA<br>GCGTAATAACAATATTATTCTGGAGTACAAGAAGCAGGATATTCTTT<br>CTCTGAATATTCCGCATGATATTAATGGTACTGAACACAGTACGCAG<br>AAGATTCAGTTGATCGTTAAGAGCAAATACGGTCTGGATCGTATCGT<br>CTGGGATGATAGTGCATTACGCAGTCAGGGCGGTCAGATTCAGCATA<br>GCGGAAGCCAAAGCGCACAAGACTACCAGGCTATTTTGCCTGCTTAT<br>GTGCAAGGTGGCAGCAATATTTATAAAGTGACGGCTCGCGCCTATGA<br>CCGTAATGGCAATAGCTCTAACAATGTACAGCTTACTATTACCGTTC<br>TGTCGAATGGTCAAGTTGTCGACCAGGTTGGGGTAACGGACTTTACG<br>GCGGATAAGACTTCGGCTAAAGCGGATAACGCCGATACCATTACTTA<br>TACCGCGACGGTGAAAAAGAATGGGGTAGCTCAGGCTAATGTCCCTG<br>TTTCATTTAATATTGTTTCAGGAACTGCAACTCTTGGGGCAAATAGT<br>GCCAAAACGGATGCTAACGGTAAGGCAACCGTAACGTTGAAGTCGAG<br>TACGCCAGGACAGGTCGTCGTGTCTGCTAAAACCGCGGAGATGACTT<br>CAGCACTTAATGCCAGTGCGGTTATATTTTTTGATGGTGCGACTAGA<br>GGTTCAAGCTCTTCGGGCTAGTCATCTAGCTCTTAA | RIVWDDSALRSQGGQIQHSGSQS<br>AQDYQAILPAYVQGGSNIYKVTA<br>RAYDRNGNSSNNVQLTITVLSNG<br>QVVDQVGVTDFTADKTSAKADNA<br>DTITYTATVKKNGVAQANVPVSF<br>NIVSGTATLGANSAKTDANGKAT<br>VTLKSSTPGQVVVSAKTAEMTSA<br>LNASAVIFFDGATRGSSSSGXSS<br>SS |
| OmpX-TAG | ATGAAAAAGATTGCATGTCTGAGCGCACTGGCAGCAGTTCTGGCATT<br>TACCGCAGGCACCAGCGTTGCAGCAACCAGCACCGTTACCGGTGGTT<br>ATGCACAGAGTGATGCACAGGGTCAGATGAATAAAATGGGTGGCTTT<br>AATCTGAAATATCGCTACGAAGAAGATAATAGTCCGCTGGGTGTTAT<br>TGGTAGCTTTACCTATACCGAAAAATCAAGAACCGCAAGCGCGGCCG<br>CAGCGTAGGCAGCCAGCGGTGATTATAACAAAAATCAGTATTATGGC<br>ATTACAGCCGGTCCGGCATATCGTATTAATGATTGGGCAAGCATTTA<br>TGGTGTTGTTGGTGTTGGTTATGGCAAATTTCAGACCACCGAATATC<br>CGACCTATAAACATGATACCAGCGATTATGGTTTTAGCTATGGTGCA<br>GGTCTGCAGTTTAATCCGATGGAAAATGTTGCACTGGATTTCAGCTA<br>TGAACAGAGCCGTATTCGTAGCGTTGATGTTGGCACCTGGATTGCCG<br>GTGTGGGTTATCGTTTTCACCATCACCATCACCATTAA | MKKIACLSALAAVLAFTAGTSVA<br>ATSTVTGGYAQSDAQGQMNKMGG<br>FNLKYRYEEDNSPLGVIGSFTYT<br>EKSRTASAAAAXAASGDYNKNQY<br>YGITAGPAYRINDWASIYGVVGV<br>GYGKFQTTEYPTYKHDTSDYGFS<br>YGAGLQFNPMENVALDFSYEQSR<br>IRSVDVGTWIAGVGYRFHHHHHH |
| PelB-NB-<br>F67TAG | ATGAAATACCTATTGCCTACGGCAGCCGCTGGATTGTTATTACTCGC<br>GGCCCAGCCGGCCATGGCCGGTTCTGAAGTTCAGCTGCTGGAATCTG<br>GTGGTGGTCTGGTTCAGGCGGGTGACTCTCTGCGTCTGTCTTGCGCG<br>GCGTCTGGTCGTACCTTCTCTGCGTACGCGATGGGTTGGTTCCGTCA<br>GGCGCCGGGTAAAGAACGTGAATTCGTTGCGGCGATCTCTTGGTCTG<br>GTAACTCTACCTACTACGCGGACTCTGTTAAAGGTCGTTAGACCATC<br>TCTCGTGACAACGCGAAAAACACCGTTTACCTGCAGATGAACTCTCT<br>GAAACCGGAGGACACCGCGATCTACTACTGCGCGGCGCGTAAACCGA<br>TGTACCGTGTTGACATCTCTAAAGGTCAGAACTACGACTACTGGGGT<br>CAGGGTACCCAGGTTACCGTTTCTTCTCACCACCACCACCACCACTA<br>A | MKYLLPTAAAGLLLLAAQPAMAG<br>SEVQLLESGGGLVQAGDSLRLSC<br>AASGRTFSAYAMGWFRQAPGKER<br>EFVAAISWSGNSTYYADSVKGRX<br>TISRDNAKNTVYLQMNSLKPEDT<br>AIYYCAARKPMYRVDISKGQNYD<br>YWGQGTQVTVSSHHHHHH |
| MBP wild-type<br>(W10/W62/<br>W94/W128/<br>W158/W230/<br>W232/W340<br>TAG)[b,c] | ATGAAACATCACCATCACCATCACCCCATGAGCGATTACGACATCCC<br>CACTACTGAGAATCTTTATTTTCAGGGCCATATGAAAATCGAAGAAG<br>GTAAACTGGTAATCTGGATTAACGGCGATAAAGGCTATAACGGTCTC<br>GCTGAAGTCGGTAAGAAATTCGAGAAAGATACCGGAATTAAAGTCAC<br>CGTTGAGCATCCGGATAAACTGGAAGAGAAATTCCCACAGGTTGCGG<br>CAACTGGCGATGGCCCTGACATTATCTTCTGGGCACACGACCGCTTT<br>GGTGGCTACGCTCAATCTGGCCTGTTGGCTGAAATCACCCCGGACAA<br>AGCGTTCCAGGACAAGCTGTATCCGTTTACCTGGGATGCCGTACGTT<br>ACAACGGCAAGCTGATTGCTTACCCGATCGCTGTTGAAGCGTTATCG<br>CTGATTTATAACAAAGATCTGCTGCCGAACCCGCCAAAAACCTGGGA<br>AGAGATCCCGGCGCTGGATAAAGAACTGAAAGCGAAAGGTAAGAGCG<br>CGCTGATGTTCAACCTGCAAGAACCGTACTTCACCTGGCCGCTGATT<br>GCTGCTGACGGGGGTTATGCGTTCAAGTATGAAAACGGCAAGTACGA<br>CATTAAAGACGTGGGCGTGGATAACGCTGGCGCGAAAGCGGGTCTGA<br>CCTTCCTGGTTGACCTGATTAAAAACAAACACATGAATGCAGACACC<br>GATTACTCCATCGCAGAAGCTGCCTTTAATAAAGGCGAAACAGCGAT<br>GACCATCAACGGCCCGTGGGCATGGTCCAACATCGACACCAGCAAAG | MKHHHHHHPMSDYDIPTTENLYF<br>QGHMKIEEGKLVIXINGDKGYNG<br>LAEVGKKFEKDTGIKVTVEHPDK<br>LEEKFPQVAATGDGPDIIFXAHD<br>RFGGYAQSGLLAEITPDKAFQDK<br>LYPFTXDAVRYNGKLIAYPIAVE<br>ALSLIYNKDLLPNPPKTXEEIPA<br>LDKELKAKGKSALMFNLQEPYFT<br>XPLIAADGGYAFKYENGKYDIKD<br>VGVDNAGAKAGLTFLVDLIKNKH<br>MNADTDYSIAEAAFNKGETAMTI<br>NGPXAXSNIDTSKVNYGVTVLPT<br>FKGQPSKPFVGVLSAGINAASPN<br>KELAKEFLENYLLTDEGLEAVNK<br>DKPLGAVALKSYEEELAKDPRIA<br>ATMENAQKGEIMPNIPQMSAFXY |

| | | |
|---|---|---|
| | TGAATTATGGTGTAACGGTACTGCCGACCTTCAAGGGTCAACCATCC<br>AAACCGTTCGTTGGCGTGCTGAGCGCAGGTATTAACGCCGCCAGTCC<br>GAACAAAGAGCTGGCGAAAGAGTTCCTCGAAAACTATCTGCTGACTG<br>ATGAAGGTCTGGAAGCGGTTAATAAAGACAAACCGCTGGGTGCCGTA<br>GCGCTGAAGTCTTACGAGGAAGAGTTGGCGAAAGATCCACGTATTGC<br>CGCCACCATGGAAAACGCCCAGAAAGGTGAAATCATGCCGAACATCC<br>CGCAGATGTCCGCTTTCTGGTATGCCGTGCGTACTGCGGTGATCAAC<br>GCCGCCAGCGGTCGTCAGGCTGTCGATGAAGCCCTGAAAGATGCACA<br>GACTCGTATCACCAAGTAA | AVRTAVINAASGRQAVDEALKDA<br>QTRITK |
| hPXR-LBD wild-type (W199/W223/ W299TAG)[b] | ATGAGCGAACGCACCGGTACACAGCCGCTGGGTGTTCAGGGTCTGAC<br>CGAAGAACAGCGTATGATGATTCGTGAACTGATGGATGCACAGATGA<br>AAACCTTTGATACCACCTTTAGCCACTTCAAAAACTTTCGTCTGCCT<br>GGTGTTCTGAGCAGCGGTTGTGAACTGCCGGAAAGTCTGCAGGCACC<br>GAGCCGTGAAGAGGCAGCAAAATGGTCACAGGTTCGTAAAGATCTGT<br>GTAGCCTGAAAGTTAGCCTGCAGCTGCGTGGTGAAGATGGTAGCGTT<br>TGGAACTATAAACCGCCTGCAGATAGCGGTGGTAAAGAAATCTTTAG<br>CCTGCTGCCGCATATGGCAGATATGAGCACCTATATGTTTAAAGGCA<br>TTATCAGCTTCGCCAAGGTGATTAGCTATTTTCGTGATCTGCCGATT<br>GAAGATCAGATCAGCCTGCTGAAAGGTGCAGCATTTGAACTGTGTCA<br>GCTGCGTTTTAATACCGTGTTTAATGCAGAAACCGGCACCTGGGAAT<br>GTGGTCGTCTGAGCTATTGTCTGGAAGATACAGCCGGTGGTTTTCAG<br>CAACTGCTGCTGGAACCGATGCTGAAATTCCATTATATGCTGAAGAA<br>ACTGCAACTGCACGAAGAGGAATATGTTCTGATGCAGGCAATTAGCC<br>TGTTTAGTCCGGATCGTCCGGGTGTACTGCAGCATCGTGTTGTTGAT<br>CAGCTGCAAGAACAGTTTGCAATTACCCTGAAAAGCTATATCGAATG<br>CAATCGTCCGCAGCCTGCACATCGTTTTCTGTTTCTGAAAATTATGG<br>CAATGCTGACCGAACTGCGTAGTATTAATGCACAGCATACCCAGCGT<br>CTGCTGCGTATTCAGGATATTCATCCGTTTGCAACACCGCTGATGCA<br>AGAACTGTTCGGTATTACCGGTAGCGGTGGCAGCGGTGGTTCAAGCC<br>ATAGCAGCCTGACAGAACGTCATAAAATTCTGCACCGCCTGCTGCAA<br>GAGGGTAGCCCGAGCCATCACCATCATCATCATTAA | MSERTGTQPLGVQGLTEEQRMMI<br>RELMDAQMKTFDTTFSHFKNFRL<br>PGVLSSGCELPESLQAPSREEAA<br>KXSQVRKDLCSLKVSLQLRGEDG<br>SVXNYKPPADSGGKEIFSLLPHM<br>ADMSTYMFKGIISFAKVISYFRD<br>LPIEDQISLLKGAAFELCQLRFN<br>TVFNAETGTXECGRLSYCLEDTA<br>GGFQQLLLEPMLKFHYMLKKLQL<br>HEEEYVLMQAISLFSPDRPGVLQ<br>HRVVDQLQEQFAITLKSYIECNR<br>PQPAHRFLFLKIMAMLTELRSIN<br>AQHTQRLLRIQDIHPFATPLMQE<br>LFGITGSGGSGGSSHSSLTERHK<br>ILHRLLQEGSPSHHHHHH |

[a] X indicates the positions of CN-Trp.

[b] Red TGG codons identify the sites of tryptophan residues in the wild-type amino acid sequence, which were individually mutated to amber stop codons to produce different protein samples with single 4-CN-Trp residues.

[c] We discovered late in the project that the template used for the present work contained the mutation T356A compared to the commonly accepted amino acid sequence of wild-type *E. coli* MBP. The side chain of this residue is solvent-exposed and the local structure is practically unchanged relative to the previously published structure of wild-type MBP (PDB ID 1OMP[55]).

**Table S4.** Table of crystallographic statistics of MBP-WT and MBP-4CNW.

| **PDB ID** | 9CLD | 9CLC |
|---|---|---|
| **Data collection** | | |
| Source | Australian Synchrotron MX2 | Australian Synchrotron MX2 |
| Space group | P 1 $2_1$ 1 | P 1 $2_1$ 1 |
| Cell dimensions | | |
| *a, b, c* (Å) | 43.82 64.52 57.64 | 43.91 64.66 57.73 |
| α, β, γ (°) | 90.0 101.2 90.0 | 90.0 101.4 90.0 |
| Resolution (Å) | 42.99 – 1.58 (1.62 – 1.58) | 42.56 – 1.48 (1.51 – 1.48) |
| $R_{merge}$ | 0.074 (0.933) | 0.084 (0.851) |
| $R_{pim}$ | 0.037 (0.464) | 0.048 (0.488) |
| $I/\sigma I$ | 10.6 (1.7) | 10.0 (2.2) |
| $CC_{1/2}$ | 0.998 (0.635) | 0.995 (0.464) |
| Completeness (%) | 99.5 (99.3) | 96.2 (94.4) |
| Redundancy | 1.8 (1.8) | 4.0 (4.0) |
| **Refinement** | | |
| Resolution (Å) | 42.99 – 1.58 (1.62 – 1.58) | 42.56 – 1.48 (1.51 – 1.48) |
| No. reflections | 42956 (2845) | 50774 (2753) |
| $R_{work}/R_{free}$ | 0.1583 / 0.1763 (0.2373 / 0.2521) | 0.1538 / 0.1657 (0.2639 / 0.2962) |
| Number of atoms | | |
| Protein | 2916 | 2928 |
| Ligand/ion | 88 | 107 |
| Solvent | 316 | 298 |
| *B*-factors (overall) | 25.0 | 23.0 |
| Protein | 22.3 | 21.0 |
| Ligand/ion | 48.7 | 48.9 |
| Solvent | 43.7 | 33.8 |
| RMS deviations | | |
| Bond lengths (Å) | 0.007 | 0.006 |
| Bond angles (°) | 0.91 | 0.87 |

[a] Values in parentheses are for the highest resolution shell.
